# Thermodynamic principles link *in vitro* transcription factor affinities to single-molecule chromatin states in cells

**DOI:** 10.1101/2025.01.27.635162

**Authors:** Julia M. Schaepe, Torbjörn Fries, Benjamin R. Doughty, Olivia J. Crocker, Michaela M. Hinks, Emil Marklund, William J. Greenleaf

## Abstract

The molecular details governing transcription factor (TF) binding and the formation of accessible chromatin are not yet quantitatively understood – including how sequence context modulates affinity, how TFs search DNA, the kinetics of TF occupancy, and how motif grammars coordinate binding. To resolve these questions for a human TF, erythroid Krüppel-like factor (eKLF/KLF1), we quantitatively compare, in high throughput, *in vitro* TF binding rates and affinities with *in vivo* single molecule TF and nucleosome occupancies across engineered DNA sequences. We find that 40-fold flanking sequence effects on affinity are consistent with distal flanks tuning TF search parameters and captured by a linear energy model. Motif recognition probability, rather than time in the bound state, drives affinity changes, and *in vitro* and *in nuclei* measurements exhibit consistent, minutes-long TF residence times. Finally, pairing *in vitro* biophysical parameters with thermodynamic models accurately predicts *in vivo* single-molecule chromatin states for unseen motif grammars.

**Highlights:**

1. KLF1:DNA binding is consistent with a three state binding model wherein the probability of recognizing the motif from a nonspecifically-bound state drives affinity.
2. Substantial effects from proximal and distal motif-flanking sequence on KLF1 binding affinity are captured by an extended PWM model.
3. KLF minutes-long residence time inferred from single-molecule footprinting *in nuclei* is consistent with *in vitro* measurement.
4. *In vitro* binding energies combined with thermodynamic models predict *in vivo*, single-molecule chromatin configurations across motif grammars and sequence contexts.

## Introduction

Sequence-specific binding of transcription factors (TFs) to DNA is essential for spatiotemporal regulation of eukaryotic gene expression. TFs interact with the chromatin landscape to bind cognate motifs and establish active regulatory elements^1^. However, the molecular details that drive TF binding events and the subsequent establishment of accessible chromatin regions (often observed through bulk accessibility measurements) are not understood from first principles – including the manner by which TFs scan DNA for binding sites, how affinity is modulated by local sequence context, the on- and off-rate kinetics of TF binding, and how motif grammars (i.e. spacing and affinity) coordinate to establish TF-occupied, nucleosome-depleted regions capable of regulating gene expression.

A physical model of TF-DNA binding requires understanding how a TF searches and identifies its cognate motif on accessible DNA, and how this interaction then affects nucleosome occupancy. The classical two-state model of protein-substrate binding posits that variable binding strength across DNA sequences is governed by dissociation rate, while diffusion-limited association rates remain constant. Many have noted that this simplified view is at odds with the speed-specificity paradox in TF search^2^, neglects nonspecific diffusion along DNA^3–7^, and cannot explain observations of sequence-dependent association rates^8,9^. Recent work with the bacterial *lac* repressor^10^ found that the probability of recognizing the motif and transitioning from a nonspecific to a specific interaction, rather than the time spent in the bound state, dictated binding strength. However, the extent to which these observations are true for eukaryotic TFs, which have weaker and less specific interactions with cognate motifs than bacterial TFs, is currently not known, and resolving these molecular details would provide a quantitative microscopic model of the interactions governing TF search and motif recognition in eukaryotes.

Furthermore, measurement of eukaryotic TF binding kinetics *in vitro* and *in vivo* often demonstrates surprising discrepancies. TF residence times measured in live cells (generally via single molecule fluorescence methods that do not resolve the genomic localization of the DNA-protein interaction) are reported to be on the seconds timescale and within an order of magnitude of nonspecific DNA-protein interactions^11–17^. In contrast, the residence times measured *in vitro* to known DNA sequences are orders of magnitude longer than the nonspecific interactions^16^. Recent work has identified subpopulations of long-duration binding events *in vivo*^*18–21*^, suggesting that live cell residence times can be underestimated due to technical limitations. Non-imaging methods to quantitatively measure TF off-rates at pre-identified binding sites would provide a key orthogonal approach to dissecting binding kinetics in cells.

Beyond these kinetic discrepancies, *in vitro* derived thermodynamic parameters also fail to accurately predict *in vivo* equilibrium binding sites, with only a small fraction (<1%) of predicted motifs exhibiting appreciable TF binding^22–25^. One possible explanation of this lack of *in vitro* and *in vivo* correspondence is the substantial effects that DNA sequence context outside of the motif has on both the TF binding search process and motif recognition. Prior work has demonstrated that motif-flanking sequence can substantially tune TF occupancy through a range of mechanisms -by establishing a more favorable DNA shape at the motif^26–31^, creating a funneled energy landscape through low-affinity motifs overlapping the core motif^32^ or homotypic environments^33^, or providing opportunities to directly bind the flanks through weak but entropically favorable sites such as those in repetitive elements^34–39^. However, much of this work is *in vitro* only, specific to bHLHs and motif-proximal bases, and focused solely on equilibrium measurements rather than TF binding kinetics.

Finally, genomic enhancers are not just composed of one isolated motif and its sequence context. Rather, these elements contain a collection of motifs^40,41^, enabling the potential for cooperative interactions between TFs mediated through direct physical interactions such as DNA allostery^42,43^ or indirect mechanisms like nucleosome competition or cofactor recruitment^44– 47^. Recent work quantitatively explained the binding of a synthetic TF across increasing motif numbers with a statistical-mechanics-based thermodynamic model that invoked these indirect mechanisms of cooperativity^47^. However, other aspects of motif grammar such as spacing have been shown to drive direct TF cooperativity *in vitro*^*43*^. Pairing *in vitro* measurements across all aspects of motif grammar with integrative thermodynamic models and *in vivo* observations promises to provide a mechanistic model linking sequence features to TF and nucleosome occupancy.

To develop this physical framework capable of predicting chromatin configurations and TF binding in cells, we paired high-throughput *in vitro* biochemical measurements with *in vivo* quantification of TF and nucleosome occupancies across individual molecules. Our model TF, Krüppel-like factor 1 (KLF1/eKLF), is a C2H2 zinc finger protein (ZNF), the most common class of human TFs^48^. KLF1’s three-finger DNA-binding domain (DBD) is highly conserved across the KLF family^49^, which enacts wide-ranging roles across development^50^ with KLF1 being critical to erythropoiesis and the switch from fetal to adult hemoglobin expression^51^.

We first measure KLF1’s binding kinetics and affinity to thousands of DNA sequences *in vitro* and find that the probability of recognizing the motif, rather than the time spent in the bound state, dictates binding strength. Motif-proximal bases drive a ∼40-fold variation in affinity that is well-described by a linear energy contribution per flanking base pair and is likely driven by direct contacts from an unconventional zinc finger binding mode. Additionally, distal sequence helical turns away from the motif can tune affinity up to 2-fold, in a manner consistent with sequence-dependent TF search dynamics. We then implemented single-molecule footprinting (SMF)^52–54^ on a library of genome-integrated constructs in an erythroleukemia cell line to quantify TF and nucleosome occupancies along individual molecules. A footprinting-based measure of KLF residence time at known motifs *in nuclei* agrees with *in vitro* measurement on a timescale of minutes. We implement thermodynamic models to explain how nucleosome competition drives nonlinear KLF occupancy across motif numbers and to accurately predict *in vivo*-specific effects from altered motif spacing. Lastly, by combining our *in vivo* thermodynamics and *in vitro* flanking sequence models, we predict how sequence context tunes occupancy in cells. Taken together, our findings demonstrate that quantitative measurement of TF binding *in vitro* and *in vivo* enables a quantitative biophysical understanding of the sequence features that lead to the establishment of accessible chromatin elements.

## Results

### High-throughput measurement of KLF1 binding kinetics and affinities

To measure KLF1 affinity to DNA *in vitro*, we designed a library of 15,651 DNA sequences containing variants within the nine-base pair KLF motif. This library spans all single, all double and some triple mutations to the consensus motif (CCACGCCCA), as well as extensive programmed variation in the flanking sequence around the motif, and negative control sequences without the motif (Methods, **Fig. 1A**). We then performed ‘high-throughput sequencing’-’fluorescent ligand interaction profiling’ (HiTS-FLIP^55^) by sequencing the library on an Illumina Miseq and using the resultant flow cell to quantify DNA binding of purified and TMR-labeled KLF1 DBD (**Fig. 1B, Fig. S1A**). We assayed binding kinetics by imaging while flowing in KLF1 to measure relative association rates (*k*_a_/*k*_*a*,*ref*_) and, post-equilibration, flowing in buffer alone to measure dissociation rates (*k*_d_) (Methods, **Fig. 1C-D, Fig S1B**). The ratios of association and dissociation rates yield relative affinities for each DNA sequence (Methods) that are broadly consistent with fitted dissociation constants (*K*_D_) from equilibrium binding measurements across KLF1 concentrations (**Fig. S1C-D**). The resulting KLF1 position weight matrix (PWM) constructed from motif mutations yields a CACCC-box, the DNA element canonically recognized by the Sp/KLF family, with finger 1 contributing the least to overall affinity as has been previously observed with homologs KLF4 and Sp1^56–58^ (**Fig. 1E**).

**Figure 1:**
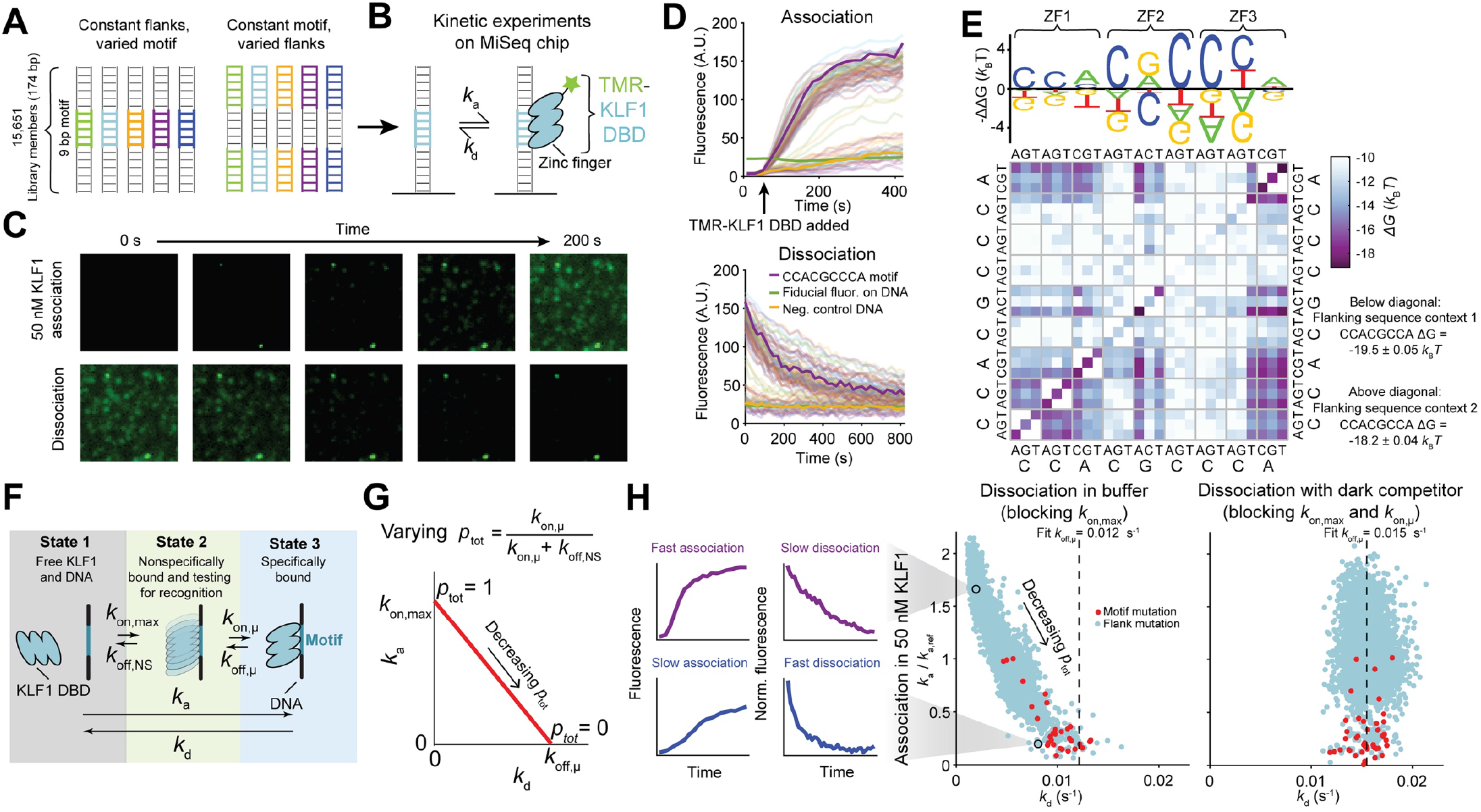
KLF1:DNA binding is driven by motif recognition probability in high-throughput *in vitro* kinetic measurements. A) Schematic of DNA library with variable KLF1 motifs and flanking sequences. B-C) Schematic (B) and images (C) of KLF1 binding to DNA targets in a sequenced flow cell. A) KLF1 association (top) and dissociation (bottom), including 100 examples across the library (faded lines). B) Binding energies (Δ*G* = ln(*K*_D_)) for KLF1 across single and double mutations to the core motif in two sequence contexts. Single mismatch mutations for sequence context 1 are on the diagonal. A position weight matrix (PWM) fitted on the motif mutants is represented on top. C) Schematic of the kinetic model describing KLF1-DNA binding. D) The effective rate constants for the association to (*k*_a_) and dissociation from (*k*_d_) the target site are coupled according to the equation^10^: *k*_a_ = *k*_on, max_ -(*k*_on, max_ / *k*_off, u_) x *k*_d_. This relationship becomes anticorrelated and linear when *k*_off,μ_ is constant and *p*_tot_ changes (red line). E) Measured association and dissociation rate constants for motif (red) and flank (blue) mutations. Dissociation is measured with (right) and without (middle) dark competitor (unlabeled high concentration KLF1). Dotted lines are fitted microscopic dissociation rates (*k*_off, u_). Insets on the left are representative associations and dissociation curves for high affinity (top, purple) and low affinity (bottom, blue) binding.

### KLF1 affinity is driven by motif recognition probability

We first tested whether our binding measurements were in agreement with a recent model^10^ that postulates a non-specific and transient DNA-associated TF “testing” state, from which the TF can either specifically bind the target sequence (*k*_on,u_) or dissociate into solution (*k*_off,NS_) (**Fig. 1F**). If affinities for different targets vary due to altered probabilities for specifically binding the motif (*p*_tot_ = *k*_on,u_/(*k*_on,u_+*k*_off,NS_)), then this three-state microscopic model predicts an anticorrelation between macroscopic association and dissociation rates (**Fig. 1G**). Indeed, we found that a coupling between association and dissociation rates, which was previously observed for the prokaryotic *lac* repressor^10^, also holds for the eukaryotic TF KLF1 across mutations to the core motif and to motif-flanking sequences (**Fig. 1H**). This three-state model predicts that if microscopic reassociation (*k*_on,u_) is blocked during dissociation (for example with a binding competitor), and if dissociation from the testing state into solution (*k*_off,NS_) is fast, the variability in macroscopic dissociation rates (*k*_d_) should converge on the microscopic dissociation rate (*k*_off,u_). Indeed, when we performed KLF1 dissociation in the presence of dark competitor (a high concentration of unlabeled KLF1), the observed dissociation rates exhibited a substantial decrease in variability, collapsing near the fit microscopic dissociation rate (**Fig 1H**). These results suggest that motif recognition probability (i.e. the transition to specific binding), rather than the time spent in the bound conformation (i.e. the residence time), drives affinity for the eukaryotic TF KLF1.

### KLF1 affinity is tuned by motif-flanking sequence

Mutations to flanking sequence (nucleotides outside of the core, 9 bp motif defined above) lead to substantial variability in KLF1 affinity (**Fig. 1H**). As we observe no measurable binding to sequences with flank mutants but no core motif (3,295 sequences, **Fig. S2A-B**), the observed variation in affinity is due to variable occupancy at the motif itself and not binding to low affinity sites in the flanks^34–39^. To determine the distance at which flanking bases can affect motif binding, we assayed binding to a core KLF motif embedded in quasi-random DNA sequences with increasing lengths of constant-sequence flanks (Methods, **Fig. 2A**). These constant sequences (containing the motif and proximal bases) were embedded either at a fixed position within variable background sequences or at variable positions within a fixed background sequence (Methods, **Fig. 2A**). As expected, variation in binding kinetics (i.e. *p*_*tot*_) is largest without any constant flanking sequence (i.e. allowing the first motif-flanking base pair to vary across sequences) and decreases as more flanking base pairs are held constant (**Fig. 2B**). The variation in dissociation constant (*K*_*D*_) across fully variable flanks is around 40-fold, decays to 2-fold by holding three flanking base pairs constant, and remains present even holding 15 bp of proximal sequence constant (**Fig. 2C**). These results indicate that KLF1 binding is tuned not only by the immediately proximal flanks (∼two base pairs) but also by more distal flanks.

**Figure 2:**
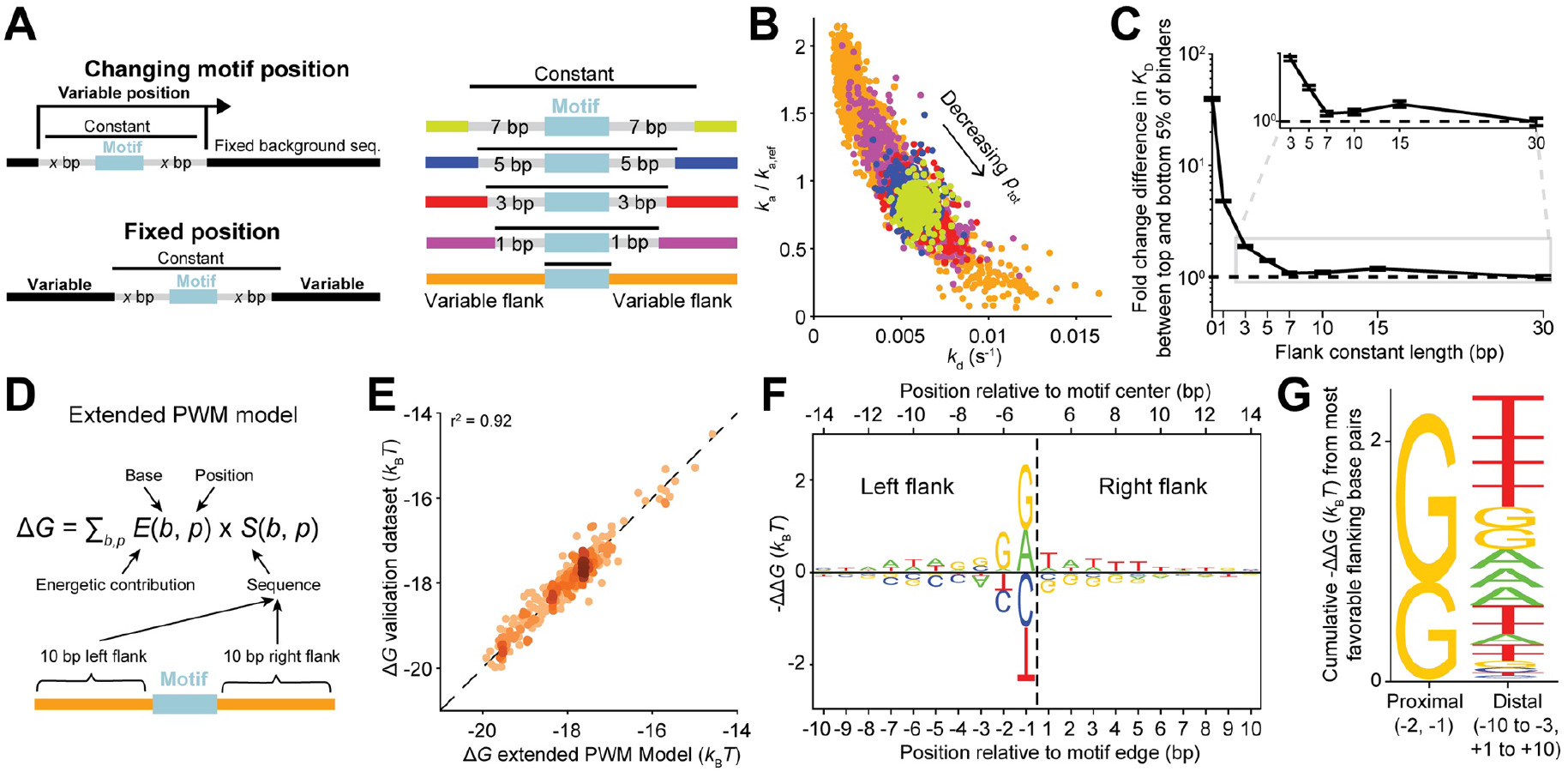
Motif-proximal flanking sequence drives 40-fold variation in KLF1 affinity. A) Schematic of libraries varying flanking sequence around the consensus KLF motif. B) Measured association and dissociation rate constants across variable lengths of constant flanks. C) Quantification of binding energy (equilibrium measurement) fold-change between top and bottom 5% of binders (set by kinetic measurement) across constant flank lengths. Error bars are 68% confidence intervals obtained by bootstrapping over clusters on the HiTS-FLIP array. D) Mathematical description of extended position weight matrix (PWM) model. E) Relationship between model predicted and measured binding energies on a held-out validation dataset. Color represents point density. F) The fit position weight matrix (PWM) represented as 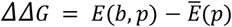,including the 10 left and right flanking base pairs. G) The cumulative contribution to binding energy for the most favorable base pairs, for the proximal flanks (positions - 2 and −1) and distal flanks (remaining positions −10 to 10).

### An extended PWM model captures proximal flanking sequence effects

To quantify the flanking sequence determinants that increase or decrease KLF1 affinity, we first fit our data with a PWM model that includes 10 flanking base pairs on each side of the core motif (**Fig. 2D**). Both the training and held-out validation data are well-fit by the extended PWM model (*r*^2^= 0.92 and 0.92, respectively; **Fig. 2E, Fig. S2C**). This extended PWM model (60 fit parameters) performed similarly to a more complex and less-interpretable machine learning model (∼1.6 million fit parameters, *r*^2^ = 0.97 and 0.95, respectively; **Fig. S2D-E**), suggesting that most variability in affinity can be accurately explained by a linear contribution to binding energy from each flanking base pair. The motif-proximal bases contribute the most to affinity, with KLF1 exhibiting a strong preference for guanines in the immediate 5’ flank (adjacent to where the first N-terminal KLF1 zinc finger binds) and a weaker preference against GC content in the 3’ flank (adjacent to where the C-terminal KLF1 zinc finger binds) (**Fig. 2F**). Consistent with this observation, many KLF1 chromatin immunoprecipitation studies identify longer motifs with guanines in the −1 or −2 positions^59–62^.

We next explored the possible mechanisms for how proximal flank effects tune affinity. First, these adjacent bases could increase affinity through creating low-affinity motifs overlapping with the core motif, as has been posited in prior studies^32^. However, when comparing the core motif PWM scores for sites overlapping the motif for low and high affinity flanking sequence groups, we find no change in the energetic landscape (**Fig. S2F**), suggesting that new low-affinity motifs do not drive the flanking sequence effects on KLF1 binding. Second, we hypothesized that proximal bases may influence the “DNA shape” at the motif to create a more favorable binding site, as has been previously suggested^26–31^. To test the explanatory power of flank-driven DNA shape effects, we used deepDNAshape^63^, a deep learning-based method that captures the influence of up to seven flanking bases, to predict key DNA shape parameters (minor groove width (MGW), propeller twist, helical twist, and roll) at the motif across the flanking sequence perturbations (**Fig. S2G**). Although we found correlations between affinity and shape parameters at the motif, their signs are inconsistent between datasets or yield negligible changes in shape parameters (**Fig. S2H-I**). These data suggest that proximal flanks do not perturb affinity through shape effects, consistent with prior findings that C2H2 zinc fingers largely rely on base rather than shape information^64^.

Lastly, proximal flanks may tune KLF1 binding through direct contacts, as interactions between C2H2 zinc fingers and adjacent base pairs outside of the canonical DNA triplet are known to occur^65^. Predicted binding modes from the structures of related homolog Sp1 reveal contacts between the first zinc finger and the phosphate backbone of proximally flanking base pairs^58^. A unique binding mode was observed in which Sp1’s first finger interacts with five base pairs, the canonical triplet and two 5’-adjacent bases^56^. Mutation of these two 5’-adjacent guanines to threonines decreases Sp1’s affinity for DNA^56^ in a manner similar to our observed proximal 5’ flank effects. This form of direct interaction between adjacent base pairs and KLF1 would be consistent with our observation that proximal flank changes can contribute more to binding energy (up to 1.4 *k*_*B*_*T*) than some bases within the core motif (down to 0.9 *k*_*B*_*T*) (**Fig. 1E, Fig. 2F**). Taken together, direct contacts driven by a unique finger one binding mode are the most likely explanation for the large proximal flank effects at positions −1 and −2.

### GC-rich sequence helical turns away from the motif decrease affinity

The two 5’ motif-adjacent bases (positions −1 and −2) dominate the variability in KLF1’s affinity. However, the combined energetic contribution to binding from the remaining flanking bases on affinity is roughly equivalent to the contribution from those two proximal bases (**Fig. 2G**) – leading to an almost 10-fold potential change in affinity. To dissect these more distal effects that are likely not driven by direct contacts, we investigated affinity to sequences where the core motif is flanked by five fixed, constant base pairs and then moved along the background sequence (**Fig. 3A**). These data show that affinity varies across positions (i.e. based on this variable distal sequence context outside of the fixed sequences). Autocorrelation analysis reveals that the affinity changes caused by the distal sequence context exhibits a periodicity of ∼10.5 base pairs, consistent with the helical pitch of B-form DNA (**Fig. 3B-C**). The amplitude of the spectral peak at the ∼10.5 bp period is largest for fixed flank lengths from three to five bps (sequence variation more than seven to nine bp away from the motif center), and decays for both longer and shorter flank constant lengths (**Fig. 3D, Fig. S3A-B**).

**Figure 3:**
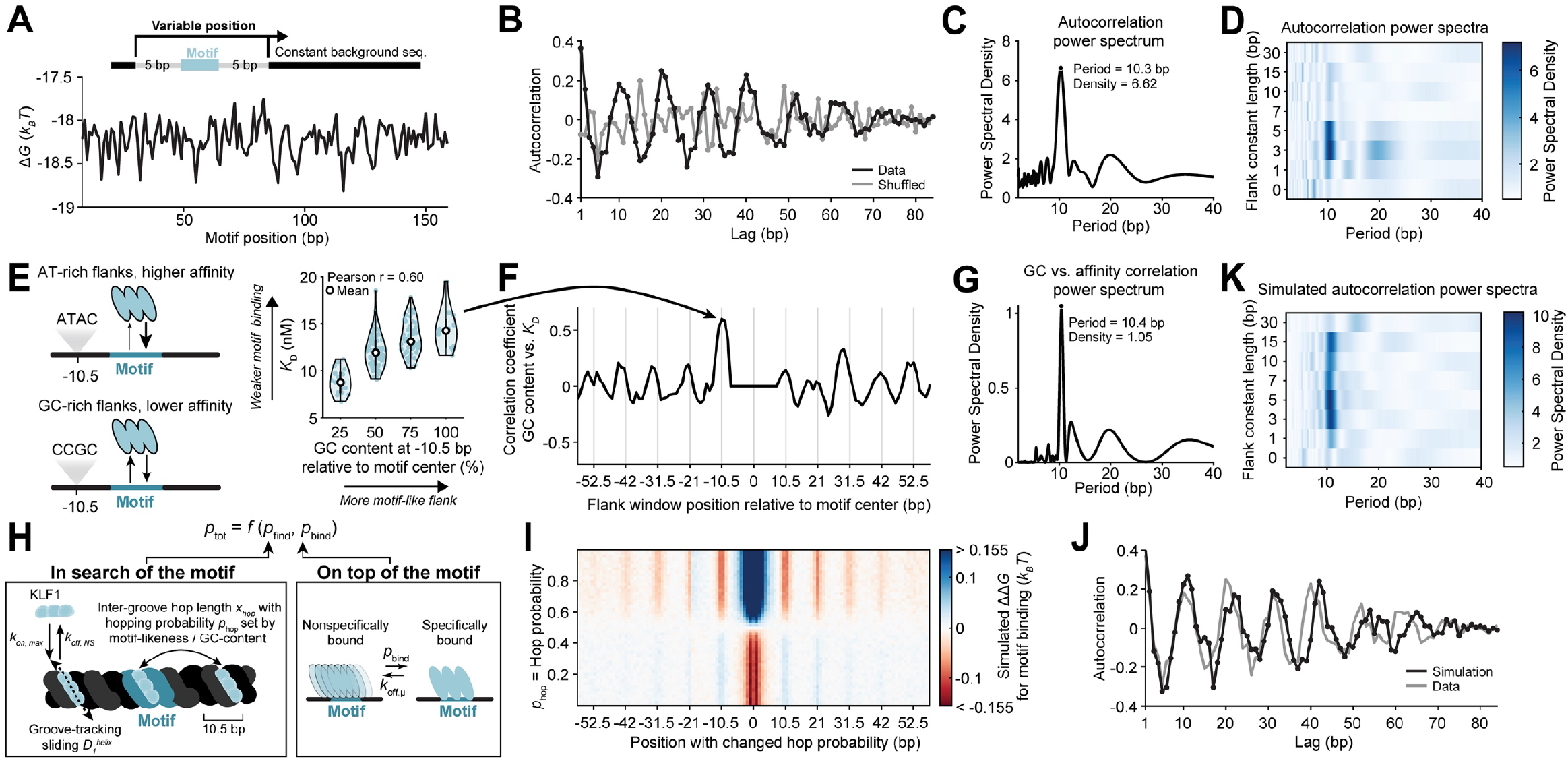
Variable KLF1 hopping probabilities in TF search explain distal sequence effects on affinity. A) Variation in binding energy across motif positions with five base pairs of constant flanks. B) Autocorrelation of the binding energies in A (black) and shuffled data (gray) across variable lag lengths. C) Power spectrum calculated via the Fourier transform of the autocorrelation in B. D) Power spectra as in C across constant flank lengths where the hue denotes the power spectral density. E) Relationship between GC content one helical turn 5’ from the motif and measured affinity. F) The correlation coefficient between GC content at variable positions away from the motif and measured affinity. G) Power spectrum of correlation coefficients in F. H) Schematic of target search in which KLF1 combined faithful DNA-groove tracking (1D diffusion) with short and frequent intergroove hops. I) Simulated relative binding energies for motif binding when the hopping probability is tuned at a single variable position relative to the motif. J) Autocorrelation of simulated binding energies where the hopping probability is set by motif-likeness of flanking sequence (black) and of measured binding energies (gray) for five base pairs of constant flanks. K) Power spectra as in D for simulated autocorrelations.

Consistent with the enrichment of AT content in favorable distal flanks (**Fig. 2G**), we observed a strong correlation between *K*_D_ and the GC content one helical turn away from the motif (Pearson r = 0.60, **Fig. 3E**). The correlation between *K*_D_ and GC content extends across the flanks with a periodicity of ∼10.5 base pairs in both the sliding and fixed position motif libraries, with the highest affinity sequences depleted in GC content helical turns away from the motif (**Fig. 3F-G, Fig. S3C-D**). Since the KLF1 motif is GC-rich, we wondered if ‘motif-like’ sequences in the flanks might be correlated with increased affinity. We used the motif PWM to score the flank sequences (see **Methods**) and observed a strong correlation between the ‘motif-likeness’ at certain positions and *K*_D_, with the degree of correlation exhibiting this periodic trend across positions as well (**Fig. S3E-G**).

### Distal flanks tuning hopping probabilities during TF search

These periodic trends cannot be explained by allosteric interactions through DNA, as has been previously observed between two TF binding sites^43,66^ for two reasons. First, negative control sequences without the core motif show no measurable binding, demonstrating that KLF1 is not specifically binding the flanks (**Fig. S2A-B**). Second, we observed the opposite correlation than would be expected from allostery, that ‘motif-like’ sequences helical turns away actually decrease measured affinity. Furthermore, we found that sequence context one helical turn away from the motif center has negligible effects on DeepDNAShape-predicted motif shape (**Fig. S2G-I**), suggesting shape preferences are not a mechanism to explain these observations.

Because the observed periodicity is close to the period of B-DNA, we then wondered if TF search dynamics might explain these distal sequence effects. TF search occurs through multiple modes: 3D diffusion, inter-segmental jumps, and both rotation-coupled (sliding) and rotation-uncoupled (hopping) 1D diffusion along DNA^4,9^. Specifically, we hypothesized that GC-rich or motif-like sequence could decrease the hopping probability, thereby reducing the likelihood of hopping one groove over onto the motif and instead increasing the likelihood of sliding along DNA and dissociating before reaching the motif. To simulate how altering the hopping landscape would impact affinity, we modified our microscopic model of TF binding (**Fig. 1F**) to break down the transition from nonspecific to specific DNA interaction into two parts: TF search for the motif along DNA and, once there, recognition of the motif. To do so, we incorporated a model for target search^9^ in which a TF combines groove-tracking sliding and hopping between major grooves (**Fig. 3H, Fig. S3H**) and then evaluated the model using stochastic reaction diffusion simulations on discretized DNA (see **Methods**). In this approach, the TF’s probability of motif recognition and transitioning from a nonspecific to a specific interaction (*p*_tot_) is deconvolved into the probability of finding the motif during 1D diffusion on DNA (*p*_find_, a function of sliding and hopping) and the probability of binding the motif when being directly on top of it (*p*_bind_) (**Fig. 3H, Fig. S3H**).

To first evaluate how the hopping landscape impacts affinity, we systematically tuned the hopping probability at individual positions. The model predicts that higher hopping probability is disfavorable adjacent to the motif and favorable at integer helical turns away from the motif (**Fig. 3I**). This trend is consistent with our experimental data if lower ‘motif-likeness’ (i.e. GC content) and weaker non-specific binding indeed lead to higher hopping probabilities. Therefore, we then used the motif-likeness across our background sequence to calculate a hopping probability at each position and directly simulated the affinity for our experimental sequences. These results recapitulate the ∼10.5 bp periodicity and decay in autocorrelation observed in our data across parameter perturbations within a physically reasonable range^9^ (**Fig. 3J,K Fig. S3I**). In addition, the model reproduces the dependence of the autocorrelation power spectra on the constant flank length (**Fig. 3D,K, Fig. S3J**). This dependence can intuitively be understood: at long constant flank lengths, effects of variable sequences very far from the motif on motif search are attenuated (**Fig. 2C**), while at very short constant flank lengths, the proximal flanking sequence effects outweigh distal effects (**Fig. 2F**). Alongside evidence against DNA allostery or DNA shape driving these distal effects (**Fig. S2A-B,G-I**), flanks tuning the hopping probability in this “goldilocks zone” during target search is the most plausible explanation for our experimental observation that sequence context perturbs the probability of transitioning from a nonspecific to specific interaction with DNA (*p*_*tot*_, **Fig. 2B**).

### Tandem *in vivo* single-molecule footprinting and *in vitro* kinetics of KLF1 binding

Given the magnitude of flanking sequence effects observed in KLF1 binding *in vitro*, we wondered if these observed *in vitro* effects drive TF occupancy and chromatin architecture differences in human cells. Methyltransferase-based single-molecule footprinting (SMF) has recently been deployed to generate high-resolution accessibility information^47,52–54^ capable of quantifying individual TF and nucleosome binding events on single molecules of engineered and genome-integrated sequences^47^. This method entails incubating isolated nuclei with methyltransferase, for example the GpC methyltransferase M.CviPI, which deposits methylation on accessible cytosines. However, the observed *in vitro* residence time for KLF1 of ∼1 minute (**Fig. 1D**) poses a substantial challenge to standard SMF workflows with over 7.5 minutes of exposure to methyltransferase. While the short residence time is a technical challenge, it also presents an opportunity to measure *in nuclei* KLF1 off-rates at cognate motifs by methylating for increasing durations of time. As KLF1 dissociates over time, we would expect its footprint of DNA protection to be “eroded” through the continuing action of methyltransferases methylating previously bound sites (**Fig. 4A**).

**Figure 4:**
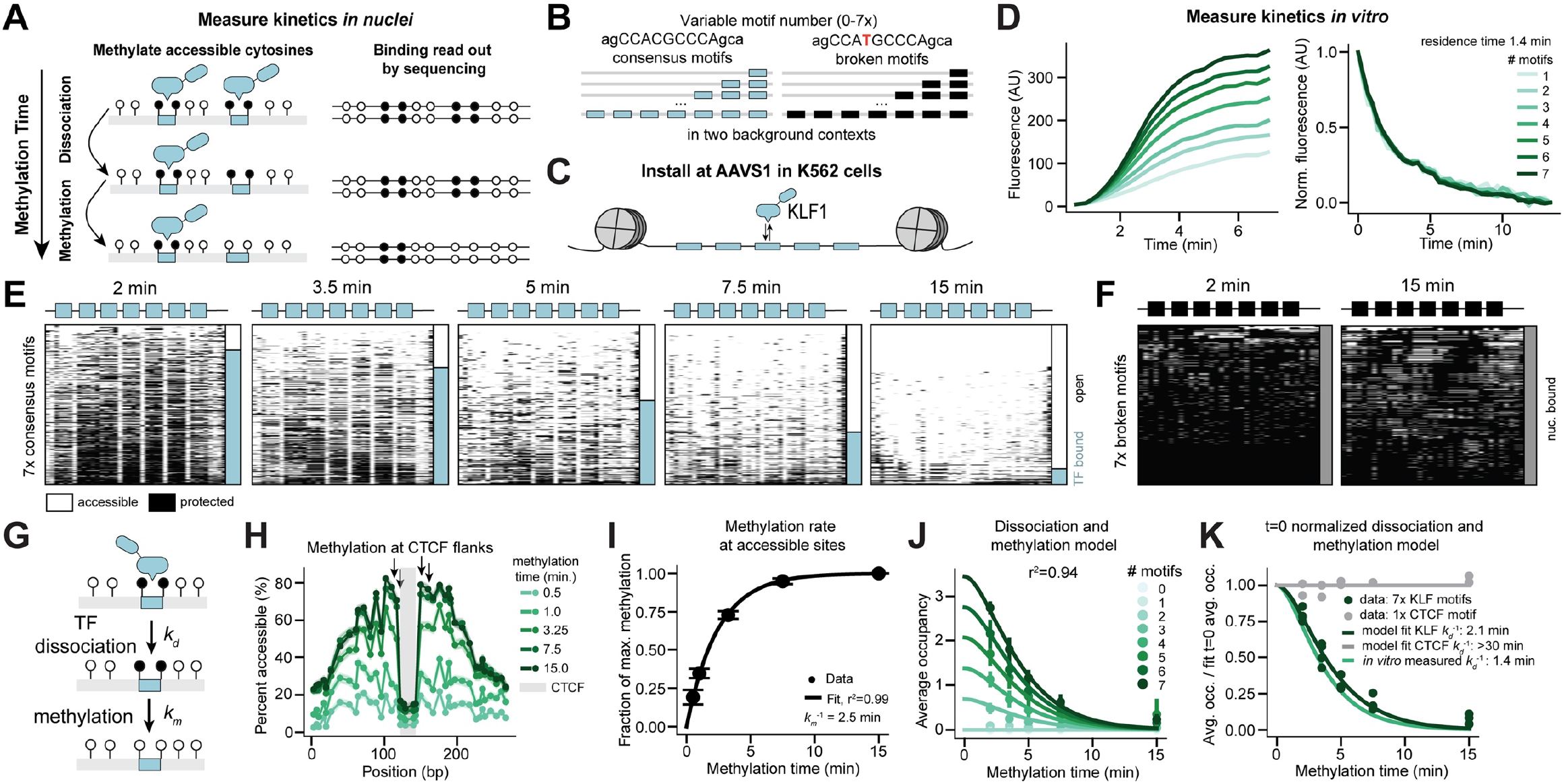
KLF residence time inferred from single-molecule footprinting in human cells is consistent with *in vitro* measurement. A) Schematic of *in nuclei* TF dissociation measurement through a methylation time course. B) Schematic of library with variable numbers for consensus (blue) and broken (black) motifs flanked by GpCs. C) Schematic of library installed at AAVS1 in K562 cells. D) *In vitro* association (left) and dissociation (right) curves for one to seven consensus motifs where dissociation is normalized to equilibrium fluorescence values. E-F) Summary plots of single molecules (rows) for molecules of constructs with seven consensus motifs from two to 15 minutes of methylation (E) or with seven broken motifs at two and 15 minutes of methylation (F). Each GpC spans an equal width in this representation; methylated Cs in white and unmethylated Cs in black. Bars show the fraction of enhancers with one or more TFs bound (blue), none bound (white), or nucleosomal (gray). F) Schematic of model for fitting TF dissociation rate from methylation time course. G) Average methylation for the CTCF construct across methylation times. Arrows represent the accessible GpCs used to fit the methylation rate. Shaded regions are bootstrapped s.d. H) Relationship between methylation time and the fraction of maximum methylation for the accessible sites denoted in H. Error bars are s.d. Line is a binding isotherm fit with a single rate. J) Relationship between methylation time and KLF occupancy across motif number. Lines are the fit dissociation and methylation model. Error bars are s.d. across two backgrounds and two biological replicates. K) Relationship between methylation time and KLF occupancy normalized to its fit occupancy at zero minute methylation time for the constructs with seven consensus KLF sites (dark green) and one CTCF site (gray) across two biological replicates. The lines are the dissociation and methylation model, fit to KLF (dark green) and CTCF (gray) data or with the input KLF off-rate from *in vitro* measurement (light green).

To measure KLF1 dissociation in cells we created a library of synthetic constructs in two sequence backgrounds containing variable numbers (0-7x) of the KLF1 consensus motif with 15 base pair spacing in between motifs (**Fig. 4B**). We flanked each motif with GCs, both to enable high-quality footprinting of KLF1 binding at these motifs with GpC methyltransferase and to maintain constant proximal flanking base pairs across binding sites. To demonstrate KLF-specific binding to this library, we included versions of these constructs where the consensus motif has a single base mutation (**Fig. 4B**) that ablates KLF1 binding more than 400-fold *in vitro* (**Fig. 1E**). We also included a construct with one high-affinity CTCF motif to serve as a control for TF dissociation, as CTCF has a longer residence time compared to other mammalian TFs in live cell measurements ^16,67–69^ and exhibits *in vitro* residence times of ∼30 minutes^70^. Furthermore, since this CTCF motif is almost always bound in cells, its flanking base pairs are highly accessible (due to statistical positioning of nearby nucleosomes^52,54,71,72^) and thus allow a measure of the M.CviPI methylation rate at accessible bases in this genomic locus.

We then integrated these constructs at the AAVS1 safe-harbor locus in K562 cells (**Fig. 4C**). K562s are an erythroleukemia cell line and KLF1 is a critical regulator for erythropoiesis^59^. RNA-seq data confirms that K562 cells have high endogenous expression of KLF1, lower levels of expression of a number of additional KLFs, and expression of a few related but less conserved SP factors (**Fig. S4A**). Because the DNA binding domain (DBD) is highly conserved across the KLF family, with almost all containing identical amino acids for the predicted DNA-contacting residues ^49^, we will refer to *in vivo* occupancy measurements as “KLF” binding.

To directly compare *in vivo* and *in vitro* binding kinetics, we first measured KLF1 associating and dissociating from these SMF-compatible constructs *in vitro*. As expected from independent binding, *in vitro* apparent association rates (the rates of appearance of fluorescence) linearly scale with motif number (**Fig. S4B**) while the *in vitro* dissociation rate is constant, with a residence time of 1.4 minutes (**Fig. 4D**). We observed no measurable association to the mutated KLF1 motif on these constructs (**Fig. S4B**).

In K562 cells, we first integrated these *in vitro*-measured constructs at AAVS1, then methylated nuclei for increasing incubation times spanning from two minutes, the shortest time at which we can achieve sufficient methylation to reliably footprint (**Fig. S4C**), to 15 minutes. DNA was then enzymatically converted^73^ to preserve methylation information during targeted amplification and Illumina sequencing of these constructs (**Fig. S4D**).

### SMF-inferred *in nuclei* KLF residence time agrees with *in vitro* measurement

In SMF data for a construct with seven consensus motifs, we observed substantial protection at motifs and accessibility in between motifs at the two minute methylation time, consistent with TF binding (**Fig. 4E, Fig. S4E**). We observe no such characteristic motif-specific protection in a construct with seven mutated motifs (**Fig. 4F, Fig. S4E**), suggesting that measured occupancy is specific to KLF binding. However, as the methylation time increases, accessibility increases across the locus and motif protection is lost (**Fig. 4E, Fig. S4E**). This loss of protection at motifs at long methylation times is consistent with KLFs dissociating from the binding site and allowing M.CviPI to methylate the motif.

To then quantify TF and nucleosome binding, we modified a probabilistic model^47^ that takes in the methylation signal and known motif locations and assigns probabilities to all configurations for each single molecule (**Fig. S4F** and Methods). To model the decrease of motif protection due to loss of KLF binding over the methylation time course, we wrote a set of ordinary differential equations (ODEs) to describe the two step process in which KLF irreversibly dissociates from its motif with rate *k*_d_ and M.CviPI then irreversibly methylates the accessible motif with rate *k*_m_ (**Fig. 4G**). While KLF could reassociate before methylation takes place, we did not include this possibility in the ODEs based on the assumption that KLF concentration greatly decreases once cells are permeabilized and resuspended in methylation buffer. We then quantified the methylation rate at this locus by fitting methylation of accessible CTCF-flanking sites with a single-rate isotherm (*r*^2^ = 0.99, *k*_m_ = 0.4 min^-1^, **Fig. 4H-I**). With this quantified methylation rate, the measurement is sensitive to quantify TF residence times longer than 30 seconds (**Fig. S4G**).

After incorporating this measured methylation rate and assuming a constant dissociation rate across motif number (as was measured *in vitro*), the occupancy data across the library and across methylation time are well fit by the dissociation and methylation ODE model (r^2^ = 0.94, **Fig. 4J**) with a fit residence time of 2.1 minutes, which is within a 2-fold margin of error of the *in vitro* measured residence time of 1.4 minutes (**Fig. 4K**). While live cell imaging typically estimates residence times on the order of seconds, a method designed to improve observation of long-term dynamics measures an *in vivo* residence time of 1.8 minutes for the closely related TF Sp1^74^, supporting our observation of a minutes-long *in vivo* residence time. Conversely, CTCF does not dissociate from its binding site throughout the methylation time course, consistent with a slow TF off-rate^67,68,70^ (**Fig. 4K, Fig. S4C**). These data demonstrate that methylation time course experiments can provide an imaging-orthogonal estimate of TF residence time in cells at known motifs that are in agreement with *in vitro* measurement on the minutes timescale.

### KLF binding exhibits cooperativity in cells but not *in vitro*

Since KLF motifs are enriched for clustered binding sites within the genome (**Fig. S5A-B**), we next investigated the relationship between motif number and KLF occupancy in cells in these data. From the probabilistic model calls of TF and nucleosome binding events on individual molecules (**Fig. 5A, Fig. S4F**), we observed that average TF occupancy is non-linear with motif number, with no observed binding with one or two motifs, and roughly linearly increasing TF occupancy from three to seven motifs (**Fig. 5B**). Nucleosome clearance demonstrates a sigmoidal relationship with motif number, with molecules completely protected until three motifs are present and completely accessible after five motifs are present (**Fig. 5C, Fig. S5C**). Consistent with KLF-specific binding, the mutated motif exhibits neither an increase in TF occupancy nor a decrease in nucleosome occupancy across motif number (**Fig. 5B-C**). Thelow-end thresholding in KLF occupancy **(Fig. 5B**) is only observed when TFs must compete with nucleosomes to bind DNA in cells, while *in vitro* binding increases linearly with motif number (**Fig. S4B**).

**Figure 5:**
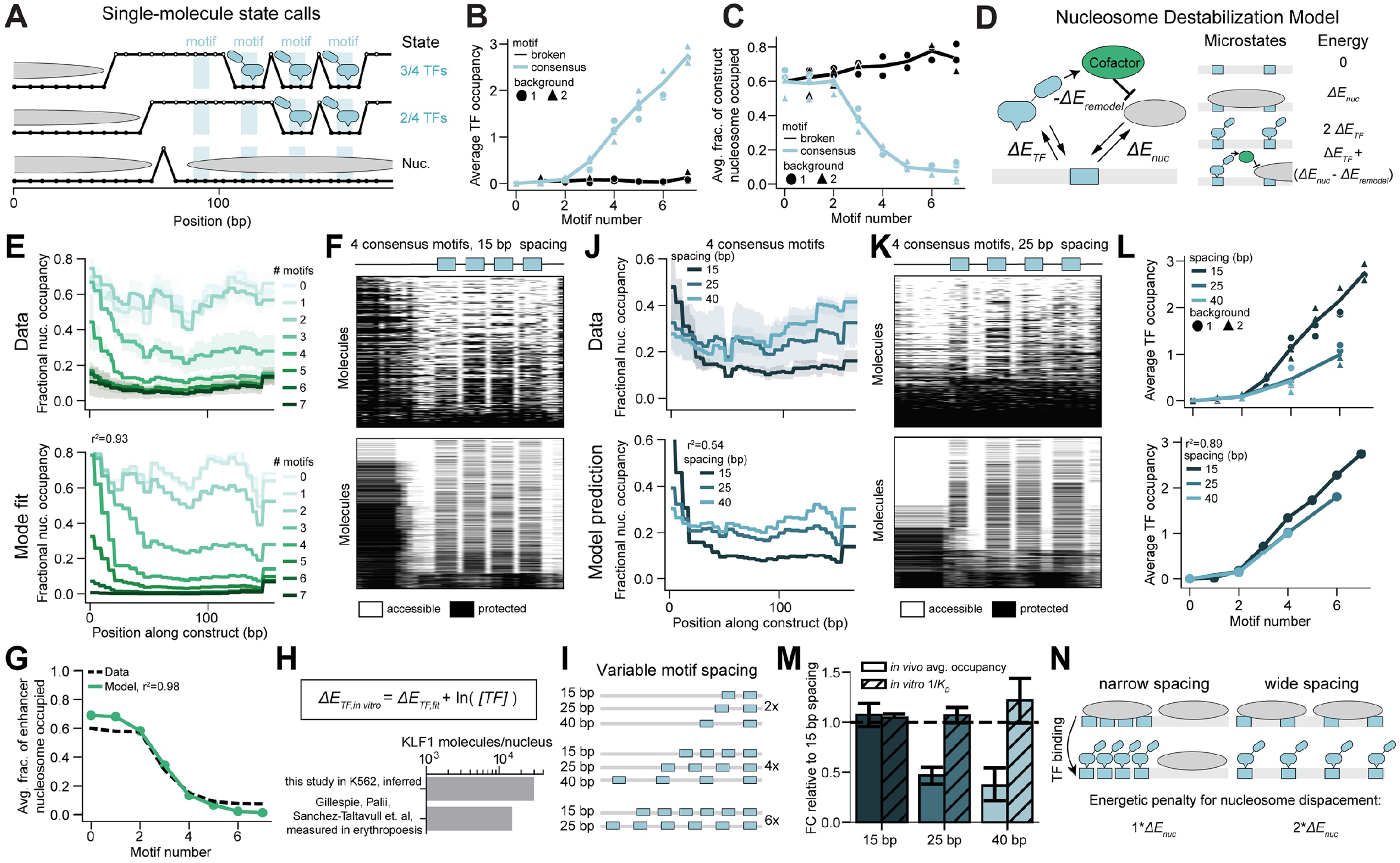
Thermodynamic models explain how nucleosome competition alters KLF binding across motif number and spacing. A) Example molecules containing four KLF consensus motifs with model-assigned molecular interpretations. Up (white) represents a methylated, accessible GpC, while down (black) represents an unmethylated, inaccessible GpC. B-C) Relationship between motif number and average KLF occupancy (B, 2 minute methylation) or the average fraction of constructs that are nucleosome occupied (C, 7.5 minute methylation) for consensus (blue) and broken (black) motifs across two background sequences (symbols) and two biological replicates. D) Schematic of the Nucleosome Destabilization model describing competition of nucleosomes and TFs for binding to DNA, with the TF-dependent-recruitment of a destabilizing cofactor. E) The fractional nucleosome occupancy across the constructs with 0-7 consensus motifs for measured (top, 7.5 minutes of methylation) and the fit Nucleosome Destabilization model (bottom). Error bars are standard deviations across two backgrounds and two biological replicates. F) Summary plots of single molecules as in Fig. 4E for molecules of constructs with four consensus motifs with 15 base pair spacing for measurements (top, 2 minute methylation) and the Nucleosome Destabilization model simulations (bottom). G) Relationship between motif number and the average fraction of constructs that are nucleosome occupied for data (black) and fit Nucleosome Destabilization model (green). H) Inferred free KLF1 molecules per nucleus from best-fit parameters for KLF binding and measured absolute KLF1 molecules per cell from^75^. I) Schematic of library with variable numbers for consensus motifs (two, four, and six) with different spacings (15, 25, and 40 bp). J) The fractional nucleosome occupancy across the constructs with four consensus motifs and variable spacing for measurements (top, 7.5 minute methylation) and Nucleosome Destabilization model predictions (bottom). Error bars are standard deviations across two backgrounds and two biological replicates. K) Summary plots as in F with 25 base pair motif spacing. L) The relationship between motif number and TF occupancy over motif spacings for measurements across two background contexts (symbols) and two biological replicates (top, 2 minute methylation) and Nucleosome Destabilization model predictions (bottom). M) Relationship between motif spacing and both *in vivo* occupancy and *in vitro* affinity compared relative to average values at 15 bp spacing. Error bars are standard deviations across two backgrounds and two biological replicates. N) Schematic of nucleosome competition when motifs are spaced close together or far apart.

### Thermodynamic model explains *in vivo* KLF binding across motif number

Thermodynamic partition functions^76–78^ have proved a useful framework for modeling non-equilibrium steady-state TF and nucleosome binding on a single-molecule level^47^. In these models, energies are assigned to both TF-DNA and nucleosome-DNA binding, and, when the TF is able to recruit chromatin remodelers, a nucleosome destabilization energy term (**Fig. 5D**). These energy parameters are then fit with maximum likelihood estimation across a single-molecule dataset by enumerating all possible binding microstates for each construct, evaluating the energy of each microstate by summing over its molecular interactions, and computing their Boltzmann probabilities. However, fitting a thermodynamic model to the KLF occupancy dataset is intractable due to dissociation-induced uncertainties in absolute *in vivo* occupancies. Instead, we fit the average nucleosome occupancy distribution across the molecules in our dataset, a metric robust to TF dissociation (**Fig. S5D**).

To this end, we first fit a simple competition model involving only TF-DNA and nucleosome-DNA binding energies (**Fig. S5E**) which was unable to recapitulate the sigmoidal accessibility across motif number or the low-end thresholding in TF occupancy, as it yields linear changes in TF and nucleosome binding across motif number (**Fig. S5F-G**). Prior SMF studies^47^ observed low-end thresholding of TF binding within cells and identified recruited chromatin remodelers, such as BAF, as key drivers of these non-linearities. Indeed, a three-parameter model^47^, which includes a nucleosome destabilization term when TFs are present (**Fig. 5D**), can fit nucleosome occupancy across our constructs, recapitulate the observed single-molecule states, and capture the lack of binding at low motif numbers (**Fig. 5E-G, L, Fig. S5H**). This observation is consistent with previous work, both *in vitro* and *in vivo*, demonstrating that KLF1 interacts directly with BAF subunits^79–81^ as well as additional cofactors^82–84^ to remodel chromatin^85^.

This partition-function model is parameterized by an effective binding energy that combines both the free energy of TF-DNA binding with the free TF concentration in the cell. Using the derived relationship between our fit energy and TF concentration (see Methods), we directly compared the partition-function fit energy with the *in vitro* energy predicted by the extended PWM model to estimate the concentration of free KLF molecules in the nucleus. We estimate ∼30,000 molecules of free KLFs per nucleus, which is within an order of magnitude of many human transcription factors’ molecule counts^86^ and is comparable to a measurement of ∼15,000 KLF1 molecules per cell during erythropoiesis^75^ (**Fig. 5H**).

### Increased motif spacing lowers KLF occupancy *in vivo*, but not *in vitro*, due to competition with nucleosomes

To test our thermodynamic model’s ability to predict nucleosome occupancy across unseen motif configurations, we designed a library with increased motif spacings from 15 base pairs to 25 and 40 base pairs for sequences with two, four, and six motifs (**Fig. 5I**). The thermodynamic model predicts that increased spacing between motifs should decrease TF binding and shift nucleosome occupancy to create shallower and wider stretches of accessibility (**Fig. 5J-L**). Experimentally, we found that increased motif spacing indeed leads to these quantitatively predicted changes to both TF and nucleosome occupancy (**Fig. 5J-L**). Since these changes are only predicted to occur when TFs are competing with nucleosomes for access to DNA, we measured *in vitro* binding, where KLF1 alone is binding DNA, across motif spacings. We observed no trend between motif spacing and affinity *in vitro* (**Fig. S5I-J**), demonstrating that the decrease in KLF occupancy as motif spacing increases is due to competition with nucleosomes for DNA (**Fig. 5M**). Therefore, this decreased occupancy arises because as motifs are less dense on the DNA, there is a larger energetic penalty for the greater nucleosome displacement needed to maintain full TF binding (**Fig. 5N**).

### *In silico* experiments support relevance of sequence context effects *in vivo*

We next aimed to determine if the sequence-context effects on affinity observed *in vitro* are sufficient to tune KLF binding within the chromatin context in cells. First, we performed an *in silico* comparison between our measured *in vitro* affinities and predicted *in vivo* binding from BPNet^87^, a machine-learning model trained on KLF1 chromatin immunoprecipitation data in K562 cells to predict peak size from DNA sequence. We observed a strong correlation between *in vitro* affinities and BPNet predictions across motif mutations (Pearson r = 0.71), indicating that *in silico* models can robustly learn TF binding preferences. With flanking sequence perturbations around a consensus motif, we observe a more moderate correlation (Pearson r = 0.42) between measured affinity and predicted *in vivo* binding (**Fig. 6A, Fig. S6A-B**), qualitatively suggesting that flanking sequence affects TF binding in cells.

**Figure 6:**
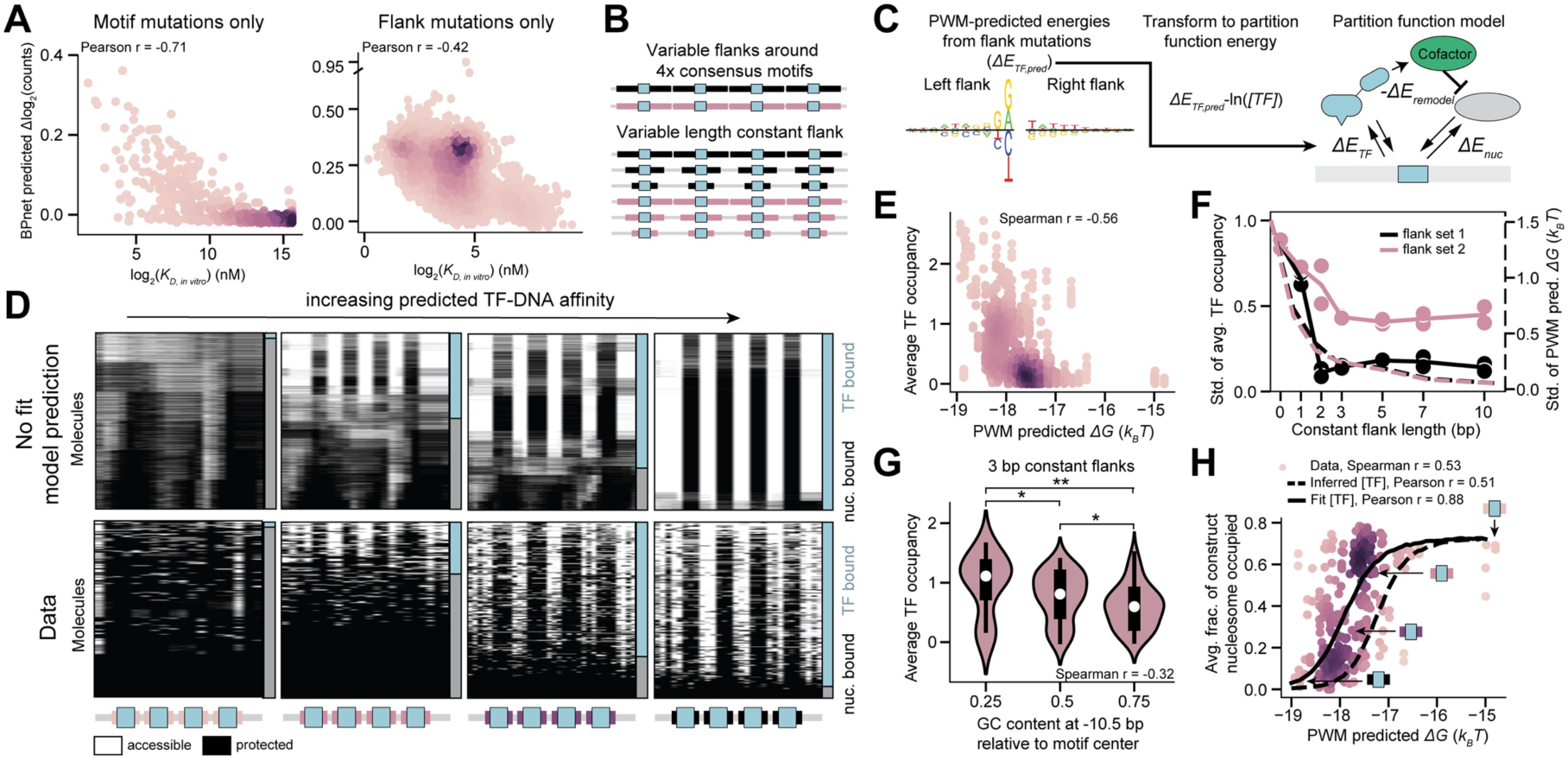
Combined extended PWM and thermodynamic models predicts *in vivo* KLF binding across motif-flanking sequences. A) Relationship between measured *in vitro* affinities and log fold-change of BPNet predicted KLF1 chromatin immunoprecipitation read counts across motif mutations (left) and flanking sequence mutations (right). Color represents point density. B) Schematic of library with four consensus KLF motifs with variable flanking sequences (top) and lengths of constant flanking sequence (bottom). C) Schematic of *in vivo* chromatin configuration predictions from the partition function model from the *in vitro* extended PWM model predicted affinities. D) Summary plots of single molecules as in Fig. 4E across constructs with four consensus motifs and variable flanks with increasingly higher affinities (columns) from the no parameter combined model predictions (top) and measurements (bottom, 2 minute methylation). E) Relationship between PWM-predicted energies and average *in vivo* TF occupancy for two biological replicates. Color represents point density. F) Relationship between the length of constant flanking sequence and the standard deviation of both *in vivo* TF average occupancy (solid lines) and *in vitro* PWM-predicted energies (dashed lines) across 15 background sequences and for two constant flank sequences (black/pink) and two biological replicates. G) Relationship between GC content one helical turn away from the motif center and the average TF occupancy across sequences from flank set two with three base pairs of constant flanks (two biological replicates). H) Relationship between PWM-predicted energies and average *in vivo* fraction of molecules that are nucleosome occupied across two biological replicates. Model predictions without fit parameters (dashed) and while fitting the TF concentration (solid) are plotted. Arrows indicate the constructs displayed in D. Color represents point density.

### Combined extended PWM and thermodynamic models accurately predict *in vivo* flanking sequence effects

We next aimed to quantitatively predict how sequence context tunes *in vivo* TF and nucleosome occupancy by combining our extended PWM model (for predicting TF energies from flanking sequence) and our thermodynamic model (for predicting binding within chromatin from TF energies). We first transform the extended PWM energies into partition function-compatible energies, enabling prediction of single molecule states from flanking sequence perturbations without any additional fit parameters (**Fig. 6C**). To then test this combined model, we designed a 300 member SMF-compatible library where each sequence contains four high-affinity consensus motifs spaced 32 bp apart with flanking sequences that are consistent across all four sites but which vary across the library (**Fig. 6B**). These variable flanks were designed to span the energetic range observed *in vitro* and to contain a sufficient number of GpCs to enable footprinting. To quantify the effect of sequence perturbations increasingly distant from the motif, we included library members with an increasing length of constant flanks for two distinct sequences and, for each constant flank length and sequence, 15 distinct background sequences for the variable region (**Fig. 6B**).

We then installed this library at AAVS1 in K562 cells and performed SMF. Despite all sequences containing these four high-affinity motifs, we observe large variability in binding as predicted by the combined models (**Fig. 6D**). Approximately 20% of the library (38 members) clearly contains non-KLF binding outside of the motif (visible in bulk accessibility plots (**Fig. S6C**)) and were therefore eliminated from further analysis. Across the remaining library members, single-molecule state calls demonstrate reproducible average TF occupancies that vary from zero to three (**Fig. 6E, Fig. S6D**). Consistent with flanking sequence perturbations driving this effect, there is a strong correlation between the extended PWM model predicted energies and *in vivo* occupancy (Spearman r = -0.56, **Fig. 6E**). To further confirm that these occupancy changes are specific to KLF binding affinity changing across sequence contexts, we performed the same experiment with transient overexpression of KLF1 (**Fig. S6G**, Methods). Indeed, occupancy is increased in mid-affinity sequences with higher KLF1 concentration, consistent with the expected effects of mass action (**Fig. S6H-I**). Taken together, these data demonstrate that bases outside of the canonical KLF motif can change the occupancy of a site from never being bound to nearly always being bound in cells.

Variability in *in vivo* KLF occupancy is largely driven by the first two flanking base pairs (which likely make direct contact with KLF1) and decreases with more constant flanking base pairs, in agreement with *in vitro* results (**Fig. 6F, Fig. S6E**). However, we observe substantial variability associated with bases more distal than these two proximal flanking base pairs (**Fig. 6F**). Therefore, we investigated if *in vivo* distal flank preferences are consistent with our finding that *in vitro* affinity is decreased by higher GC content one helical turn away from the motif. Indeed, we find that in sequences with the proximal flanking bases held constant, *in vivo* KLF occupancy decreases as distal GC content increases (**Fig. 6G, Fig. S6F**). These results demonstrate that not only do the proximal flanks’ strong effects on affinity set *in vivo* KLF occupancy, but that the more subtle effects from distal flanks, which we posit alter TF search dynamics, also contribute to tuning KLF binding in cells.

In addition to TF binding changes across the library, nucleosome occupancy (a metric robust to KLF dissociation during methylation) exhibits a sigmoidal relationship to predicted TF energies, with molecules ranging from entirely inaccessible to entirely accessible (Spearman r = 0.53, **Fig. 6H**). The combined models predict a sigmoidal response that matches the data moderately well when using the inferred TF concentration from earlier experiments (**Fig. 5H**) and well when directly fitting the TF concentration in this experiment (Pearson r = 0.51 and 0.88, respectively, **Fig. 6H**). Furthermore, transient KLF1 overexpression increased the fit TF concentration, as expected (**Fig. S6J-M**). Taken together, these data demonstrate both that the motif sequence context substantially tunes binding in cells and that biophysical models can quantitatively predict these effects.

## Discussion

Our study provides tandem *in vitro* and *in vivo* measurement of KLF1 equilibrium and kinetic binding and implements biophysical models to predict the *in vivo* occupancy of unseen sequences from *in vitro* affinities. We found that, similar to prior observations with *lac* repressor^10^, KLF1 binding strength is driven by the probability of motif recognition. This observation is consistent between these bacterial and eukaryotic TFs despite their distinct modes of DNA recognition. The dimeric *lac* repressor’s helix-turn-helix exhibits a conformational change and bends DNA upon specific binding of its 20 - 21 bp palindromic motif^88^. In contrast, monomeric zinc finger TFs like KLFs exhibit less DNA bending with conformational changes restricted to finger linker regions upon specific binding of their much shorter non-palindromic motifs^57,58^. Therefore, we expect that this microscopic model of DNA binding will likely be relevant across human TF families even though their binding mechanisms vary substantially.

The probability of motif recognition for KLF1 is dependent on both proximal and distal sequence outside of the core motif. This represents one possible explanation for how different KLF members could regulate distinct targets despite highly conserved core motifs, as has been previously observed ^60,89,90^. Since KLF1’s proximal flank preferences are likely driven by direct contacts with finger one, its ‘core motif’ likely should include bases at flank positions −2 and −1. High-throughput and quantitative measurement of flanking sequence effects on binding, such as those performed here, may reveal that core motif bases have been missed for other TFs. Furthermore, such measurements across zinc fingers and other TF families could reveal the generalizability of distal flank preferences and distinguish whether GC content (i.e. DNA stiffness) or motif-likeness drives the observed distal flank dependencies. We also anticipate that *in vitro* single-molecule experiments of target search^9^ could resolve the molecular details underlying our observed effects from distal flanks.

Our observation of a minutes-long *in nuclei* residence time for KLF contradicts the model that TF-DNA interactions are highly transient in mammalian cells^16,91,92^. Recent modifications to single-molecule tracking (SMT) enable locus-resolved observation of the yeast TF Gal4 and also report minutes-long residence times^21^, interactions which had been previously undetected by non-locus-specific SMT^93^. We anticipate that use of methods such as these, which observe TF binding kinetics at specific sites within the genome rather than across the nuclear milieu, will help to clarify the timescales of TF binding to cognate motifs, and how binding timescales relate to downstream processes such as transcriptional bursting. We note, however, that footprinting-based measurements will only quantify an upper bound of TF residence time for TFs with faster off-rates due to the time required to methylate accessible sites (**Fig. S4G**). However, more efficient or higher concentration enzymes may improve the range of residence times observable with this approach.

Here, we show that an accurate biophysical prediction of *in vivo* chromatin configurations for unseen motif grammars, such as spacing and affinity, is possible from *in vitro* derived TF binding parameters and classical thermodynamic assumptions about the nature of TF-chromatin interactions. The paired measurements both *in vitro* and *in vivo* demonstrate that inter-motif spacing can perturb TF binding through nucleosome competition without needing structural or DNA allosteric interactions as in the enhanceosome enhancer model^94^. Recently, we also demonstrated that gene expression can be explained from *in vivo* TF binding and simple physical models^47^. Together, these studies highlight that, with high-resolution and quantitative measurements, cellular processes can be understood from biophysical parameters and statistical mechanics models. We achieve accurate predictions of molecular chromatin state with relatively small numbers of physical and interpretable free parameters, in contrast to the highly parameterized “black box” deep learning models. Furthermore, we did not see evidence of highly cooperative biological state changes, such as phase separation or condensation, in contexts we observed. With the incorporation of more biophysical information into these models, such as nucleosome DNA sequence specificity and the affinities and concentrations of additional transcription factors, we anticipate that improved prediction of chromatin configurations at genomic regulatory elements will be possible.

## Methods

### Protein expression and purification

HaloTag-KLF1DBD (KLF1 DBD) was affinity purified with a N-terminal 6xHis-tag, where 6xHIS-TEV-HaloTag-KLF1DBD was expressed from a modified pD861 (ATUM) plasmid vector (pJS021) in T7-expression competent *E. coli* (NEB, C30131). 10 mL overnight cultures were diluted into 1 mL LB medium with 50 µg/mL kanamycin (Thermo Fisher, J60668.06) and grown at 37 °C and 120 RPM agitation. The expression of the protein was induced at OD_600_=0.5 by adding 0.2% weight/volume L-rhamnose (Thermo Fisher, A16166.14) and expressed for 4.5 hours. Cultures were spun down at room temperature at 5000g for 30 minutes, and frozen at −80 °C for later purification. Pellets were thawed on ice and purified using Ni-NTA Fast Start Kit (Qiagen, 30600) according to the manufacturer’s instructions. The kit lysis, wash and elution buffers were all supplemented with 1 mM DTT and one protease inhibitor tablet (Pierce, A32953). The pellet was resuspended in 10 mL lysis buffer and lysed on ice for 40 minutes. Lysate was spun down at 14,000g for 30 min and supernatant was applied to the Ni-NTA fast kit columns. The column was washed with 16 mL wash buffer (provided by kit, low imidazole concentration) and eluted with 4 mL elution buffer (provided by kit, high imidazole concentration). Eluted protein was then buffer exchanged into storage buffer (20 mM Hepes pH 7.3, 400 mM NaCl, 0.5 mM DTT) on 10 kDa cut-off Amicon Ultra-0.5 centrifugal filters (Millipore Sigma, UFC901008). Protein was then further purified with a HiTrap Heparin HP column (Cytiva, 17-5247-01). The column was first washed with 5 mL 20% ethanol and then 5 mL buffer B (20 mM Hepes pH 7.3, 2 M NaCl, 0.5 mM DTT) before equilibrating with 5 mL buffer A (20 mM Hepes pH 7.3, 250 mM NaCl 0.5 mM DTT). Protein was applied to the column, washed with 10 mL buffer A and eluted with increasing concentration of salt up to 2M NaCl in Buffer A. 6xHIS-TEV-HaloTag-KLF1DBD eluted around 775 mM NaCl and was >90% pure, evaluated via SDS-PAGE. Protein was then buffer exchanged into storage buffer on 10 kDa cut-off Amicon Ultra-0.5 centrifugal filters and diluted to 50% glycerol. Protein was then flash frozen in liquid nitrogen and stored at −80 °C for later use.

The protein was then labeled with HaloTag TMR ligand (Promega, G8251), and the His-tag was cleaved off using TEV protease (NEB, P8112S), in the same reaction. The protein was diluted to 5 µM in storage buffer supplemented with 10 µM HaloTag TMR ligand and 330 units/mL TEV protease. The reaction was incubated for 1 hour at 30 °C. Then, an additional 10 µM of HaloTag TMR was added, followed by incubation at room temperature for 2 hours. Free excess dye was then removed using Heparin purification with the same procedure as described above for the unlabeled protein. The protein was eluted with Buffer A with 775 mM NaCl. SDS-PAGE of the labeling reaction, a control labeling reaction without TEV, and collected fractions from the purification verified that excess dye did not elute with the protein and that there was a shift downwards in molecular weight expected from TEV removal of the His-tag. Protein was then buffer exchanged into storage buffer on 10 kDa cut-off Amicon Ultra-0.5 centrifugal filters and diluted to 50% glycerol. Absorbance was measured at 280 nM (A280, protein) and 548 nM (A548, TMR) with a NanoDrop spectrophotometer. The concentration of TMR in the sample was estimated using the A548 measurement, and the concentration of protein was estimated using the A280 measurement (while correcting for TMR absorbance at A280), to quantify a labeling yield of 98%. The labeled protein (TMR - KLF1 DBD) was then flash frozen in liquid nitrogen and stored at −80 °C for later use.

### *In vitro* library design

A total of 22,517 sequences were designed for *in vitro* experiments. The vast majority of these (including all results in **Fig. 1-3**) contains sequence variations in a 174 bp long variable region, varying the motif sequence, the motif position, the flanking sequence context around a motif, the number of motifs and motif spacing, and combinations thereof. Background sequences were generated randomly and any sub-sequences that resembled the KLF motif were removed. The flanking sequence context around a motif was varied both by directly mutating the flank for a constant motif position, and by changing the motif position (with varying length and identity of constant flank sequence proximal to the motif) in a constant sequence background. For the fixed motif position, flanks were mutated in a number of different ways, including randomizing the sequence 60 bp to the left and/or right of a motif (with varying length and identity of constant flank sequence proximal to the motif) and point mutating smaller regions in the 15 bp closest to the motif to the left and/or right (with varying length and identity of constant flank sequence proximal to the motif). Many of these flanking sequence context variations were also included in sequences where the strong consensus motif (CCACGCCCA) had been mutated to a dead motif (TACTTAGAT) (**Fig. S2A,B**, 3,295 sequences measured).

### *In vitro* library preparation and sequencing

The designed library variants were ordered as oligo pools from Twist Biosciences (South San Francisco, CA). The variable region of each variant was flanked by a common 5’ (ACACTCTTTCCCTACACGACGCTCTTCCGATCT) and 3’ (AGATCGGAAGAGCGGTTCAGCAGGAATGCCGAGACCG) priming sequence, enabling assembly of sequences flanked by Illumina adaptors via PCR. Libraries were prepared for sequencing by performing PCR with primers P5-TruSeqR1 and P7-TruSeqR2 (**Table S3**) and using NEBNext Ultra Q5 Master Mix (NEB, M0544) according to the manufacturer’s instructions, with 1 −10 ng starting template and 10-14 cycles of amplification. PCR reactions were then purified using AMPure XP Beads (Beckman Coulter, A63880) and DNA concentration was measured using a Qubit fluorometer. A fiducial sequence for later use in image registration was ordered as an oligo from IDT and prepared for sequencing in the same manner as for the oligo pools. Libraries were pooled with 50 - 80% PhiX Sequencing Control v3 (Illumina, FC-110-3001) and 1% fiducial sequence, and sequenced with 250 cycles in read 1 and 250 cycles in read 2 with a MiSeq Reagent Kit v3 (600-cycle) (Ilumina, MS-102-3003) on a MiSeq. Sequences and flow cell positions that completely matched a target library or fiducial sequence in read 1 were used in downstream data processing.

### HiTS-FLIP measurements

High-throughput biophysical binding measurements were performed using the sequenced MiSeq flow cell and a custom prism total internal reflection fluorescence microscope described in^95,96^. Briefly, the XY-stage, Z-stage, lasers, syringe pump, objective and camera were salvaged from an Illumina GAIIx instrument, and integrated with a fluidics adaptor that was designed to interface with Illumina MiSeq flow cells. The instrument was further equipped with a temperature control system and laser control electronics. Imaging of TMR - KLF1 DBD and TMR fiducial marker fluorescence was performed with a 532 nm laser and a 590 nm center wavelength and 104 nm guaranteed minimum 93% bandwidth bandpass emission filter (Semrock).

Following sequencing, MiSeq flow cells were prepared for binding experiments. First, DNA not covalently attached to the flow cell surface was removed by washing the flow cell with 100% formamide at 55 °C. To remove residual fluorescence from the sequencing, the flow cell was then washed with a cleavage buffer (100 mM Tris-HCl, pH 8, 100 mM NaCl, 0.05% Tween20, 100 mM TCEP) for 15 minutes at 60 °C. Fiducial marker oligo (**Table S3**) and a primer for double stranding (**Table S3**) were then hybridized to the single stranded DNA on the flow cell surface in a number of steps. First, the flow cell was washed with hybridization buffer (5x SSC buffer (ThermoFisher 15557036), 5 mM EDTA, 0.01% (v/v) Tween 20) at 60 °C. Fiducial marker (500 nM) and double stranding primer (500 nM) were then introduced into the flow cell in hybridization buffer at 60 °C and incubated for 12 minutes. The temperature was then decreased to 40 °C, followed by incubation for another 12 minutes. The flow cell was then washed with annealing buffer (1x SSC, 5 mM EDTA, 0.01% (v/v) Tween 20) at 40 °C. Fiducial marker (500 nM) and double stranding primer (500 nM) were then introduced into the flow cell in annealing buffer at 40 °C and incubated for 8 minutes. Double stranding of the DNA was then performed using the DNA polymerase I Klenow fragment (NEB, M0210S). After washing the flow cell with 1 x NEBuffer 2 (NEB, M0210S) at 37 °C, the flow cell was incubated with 0.2 U/μl Klenow fragment in 1 x NEBuffer 2 fragment for 30 minutes 37 °C. Following the double stranding reaction, the flow cell was washed with hybridization buffer.

All TMR - KLF1 DBD binding measurements were performed in an imaging buffer (10 mM Hepes, pH 7.3, 185 mM NaCl, 0.5 mg/ml BSA, 1 mM EDTA, 5 mM MgCl_2_, 0.1 mM ZnCl_2_, 0.5 mM DTT) at 22 °C. The imaging buffer was degassed using a desiccator and vacuum before each experiment. Prior to introduction of TMR - KLF1 DBD, imaging buffer was introduced into the flow cell, and fiducial markers were then imaged using a 600 ms exposure time for registration to the sequencing data.

For kinetic experiments, four tiles of the flow cells were looped over and imaged continuously, resulting in an effective imaging period over all tiles of ∼20 s. For association experiments, 50 nM TMR - KLF1 DBD was introduced into the flow cell with a flow rate of 60 μl/min during the entire experiment. After the association experiment, the dissociation experiment was performed by flowing in imaging buffer without TMR - KLF1 DBD. For the dissociation experiment with dark competitor, the imaging buffer was supplemented with 500 nM unlabeled KLF1 DBD. A flow rate of 100 μl/min was used during the entire dissociation experiments. An exposure time of 200 ms and laser fiber input power of 145 mW were used in the first association experiment and dissociation experiment, and an exposure time of 250 ms and fiber input power of 120 mW was used during the second association experiment, and dissociation experiment with dark competitor.

For the protein titration experiment, increasing concentrations of TMR - KFL1 DBD were introduced in the flow cell with a constant flow rate of 15 μl/min during the entire incubation time. For each concentration point, 14 tiles were imaged with an exposure time of 600 ms and laser fiber input power of 120 mW. The flow cell was incubated with 1.85, 5.56, 16.7, 50 and 150 Nm TMR - KFL1 DBD for 49, 49, 29, 30 and 30 minutes, respectively.

### Image processing of HiTS-FLIP measurements

For each experiment, the sequencing data were aligned with images captured during the binding experiment. Sequenced clusters were matched to fiducial marker images via an iterative cross-correlation method, following established protocols^95,97^. To account for small drifts in position during each experiment, each individual image was also aligned to the image of fiducial markers taken right before TMR - KLF1 DBD flow in. Post-alignment, fluorescence intensity for each cluster was quantified by fitting it to a 2D Gaussian.

### Fitting of kinetic and equilibrium constants

Following quantification of the fluorescence signal for each cluster, the fluorescence signal for each library member (at each time point or TMR-KLF1 DBD concentration) was calculated by taking a trimmed mean over the clusters’ fluorescence values, removing the 10% lowest and 10% highest clusters. A small subset of clusters with quantified fluorescence intensity identical to zero at all time points or TMR - KLF1 DBD concentrations (indicating unsuccessful alignment and/or Gaussian fitting of this cluster) were removed before taking the trimmed mean. This raw quantified fluorescence signal is plotted in **Fig. 1D**. This fluorescence signal was then subtracted with the fluorescence signal of a negative control sequence without KLF1 motifs (Neg. control DNA in **Fig. 1D**, seq3 in **Table S1**) to create a background subtracted fluorescence signal (**Fig. S1B top, S1C**). For dissociation experiments, a normalized fluorescence signal (**Fig. S1B bottom**) was also calculated, by dividing the background subtracted fluorescence signal with the background subtracted fluorescence signal at the first time point of dissociation for each library member. Dissociation rates constants (*k*_d_) for each library member were estimated by fitting of the equation

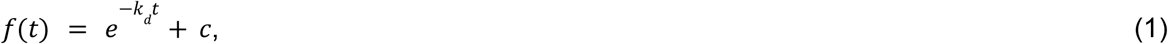

with fit parameters *k*_d_ and *c*, by minimizing the mean squared deviation of the model and the normalized fluorescence signal data values. Relative association rate constants (*k*_a_/*k*_a,ref_) were estimated via the initial slopes of the background subtracted association curves, right after TMR - KLF1 DBD flow in when the curves are approximately linear. The initial slopes were fit via linear regression by minimizing the mean squared error between the line and the data (**Fig. S1B, top**). The reference sequence (**Fig. S1B, middle**) defined to have *k*_a_/*k*_a,ref_ = 1, with reported association rate constants for all other sequences being relative to this one, is a sequence with the consensus CCACGCCCA motif and an intermediately strong flank.

Equilibrium constants (*K*_D_), from the TMR - KLF1 DBD titration experiment, were estimated using the Hill equation

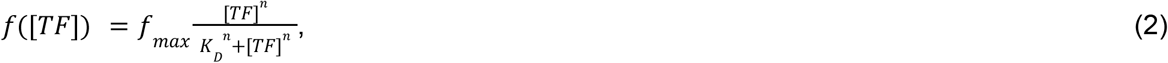

where [*TF*] is the TMR - KLF1 DBD concentration, *n* is the Hill coefficient, and *f*_max_ is the fluorescence signal at saturating TMR - KLF1 DBD occupancy at the DNA binding site. More specifically, the Hill equation was used within an algorithm which identified the most informative concentration data points for model fitting for each library member (e.g. after the fluorescence signal can be detected but before the signal saturates). We will first focus on how *K*_D_ was estimated for 0 and 1 motif sequences (all of **Fig. 1-3**), and then clarify how this procedure was modified for multiple motif sequences (**Fig. S5I**). The background subtracted fluorescence signal for the different concentration points (titration curve) was used for fitting for each library member, and, unless stated otherwise, fitting was done by minimizing the mean squared error between equation (2) and the titration curve.

First, a selection of strong binding sequences with a single consensus CCACGCCA motif, which demonstrated saturation in binding for increasing TF concentrations, were identified by selecting the sequences where the second to last concentration point had a signal that was 95% of or higher than the signal at the last concentration point. The signal at the last concentration point for these sequences was then used to create a range of *f*_max_ allowed in fitting, defined as the average of this signal ± 4 standard deviations (here quantified to be 53 ± 4×6 A.U). Equation (2) was then fit to the titration curve for each library member, while using all three of *f*_max_, *n* and *K*_D_ as free fit parameters. A global *f*_max_ was then locked to be the mode of a histogram with 20 bins (quantified to 57 A.U.) over the individually fitted *f*_max_s. To obtain an initial crude estimate of the equilibrium constant (*K*_D,initial_) for each library member, equation (2) was fit to the titration curve while locking to the global f_max_ and *n* = 1. For relatively weak binders (defined as *K*_D,initial_ < 75 nM), since the titration curve is uninformative of *f*_max_ when the curve is far from saturing, the global fit (rather than individually fit) *f*_max_ was used in later fitting.

*K*_D_ estimates were then obtained by inputting *f*_max_, *n* = 1, [*TF*], and the experimental *f*([*TF*]) into equation (2) and solving for *K*_D_, while trying to choose a [*TF*] point that had substantial fluorescence but was not close to saturation. This was implemented by choosing the 2nd titration point ([TF] = 5.56 nM) if *f*/*f*_max_>0.1, otherwise choosing the 3rd titration point if *f*/*f*_max_>0.2, otherwise choosing the 4th titration point if *f*/*f*_max_>0.2, and otherwise choosing the 5th titration point. To mitigate systematic bias based on what [*TF*] point is chosen for a given library member (which could be caused by uncertainty in [*TF*]), a correction factor was calculated for the 3rd - 5th titration points from library members that had informative (substantial binding but not saturated) fluorescence signals at multiple titration points. More specifically, the correction for a library member using the *i*th titration point was applied according to

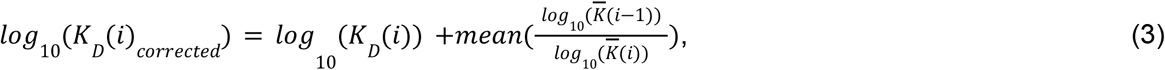

where 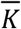 is a set of *K*_D_ values with informative fluorescence signals at both concentration *i* and *I* −1.

Initial attempts with more simple curve fitting showed that those procedures were problematic to apply unsupervised to large amounts of titration curves, since they were prone to overfit to uninformative parts of our data, either when titration curves were very low or very high and close to saturated. The described procedure does not suffer from these limitations, and should by design produce robust estimates of *K*_D_, and for ΔΔ*G* even if the uncertainty in [*TF*] is high. The excellent agreement between ΔΔ*G* measured by kinetic and titration experiments (**Fig. S1D**), validates that the fitting algorithm performs well on the current dataset.

*K*_D_ for multiple motif sequences (**Fig. S5I**) were estimated in the same way as for the 0 or 1 motif sequences, with the exception of allowing *f*_max_ to vary and be fitted over an *m* times larger interval, where *m* is the number of motifs for a given sequence. At the last step of analysis, *K*_D_ values for multiple motif sequences were divided by *m*. With this normalization performed, the occupancy of a sequence with any number of motifs (which can be greater than 1 for multiple motif sequences) will be proportional to 1/*K*_D_ at low TF concentrations, which we think is consistent and logical definition of overall *K*_D_ for DNA sequences with multiple motifs.

Binding measurements to each library member were obtained over many replicate clusters (mode of the number of clusters per library member: 22, 18, and 49 for the normal kinetic experiment, kinetic experiments where dissociation was performed with dark competitor, and protein titration experiment, respectively), enabling error estimation by bootstrapping over the clusters, with standard errors estimated as the half-width of 68% confidence intervals for *k*_a_/*k*_a,ref_, *k*_d_ and *K*_D_. For the fluorescence signals in **Fig. S1C**, standard errors were estimated as the standard deviation of the bootstrapped distribution. In **Fig. S1D**, weak binders (grey points) are defined to have *K*_D_ > 500 nM (from the titration experiment) and/or standard error in *k*_d_ > 0.0025 s^-1^. As these weak binders give noisy and uninformative *k*_d_ estimates, they were omitted from the plot and analysis in **Fig. 1H**.

### Kinetic three-state model describing TF-DNA binding

This model for TF-DNA binding (**Fig. 1F**) is a three-state markov chain, where the states are (1) free and dissociated TF and DNA, (2) a testing state where the TF is bound nonspecifically to DNA nearby or at the motif and (3) a specifically bound state where the TF is in the bound conformation on the motif. [*TF*] is the free TF concentration, *k*_*on*,*max*_[*TF*] is the association rate from solution to the nonspecifically DNA bound testing state, *k*_*off*,*NS*_ is the dissociation rate from the testing state to solution, *k*_*on*,μ_ is the rate of going from the testing state to the specifically bound conformation on the motif, and *k*_*off*,μ_ is the rate of going from the specifically bound state on the motif to the testing state. With the key assumption that the average time spent in the testing state (1/(*k*_off,NS_+*k*_on,μ_)) is very short compared to the average time spent in the free state (1/*k*_on,max_[*TF*]) and in the specifically bound state (1/*k*_off,μ_), we have previously shown^10^ that the macroscopic observables *k*_a_, *k*_d_ and *K*_D_ relate to the microscopic parameters of the model according to

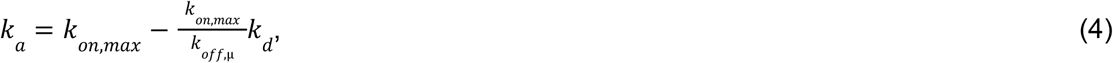

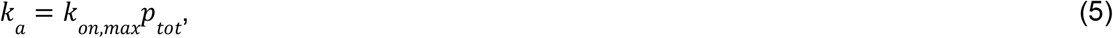

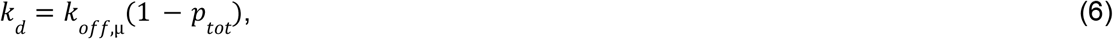

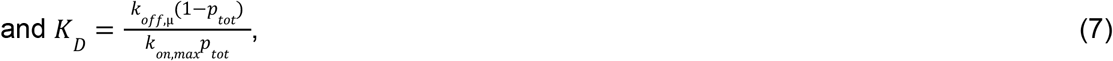

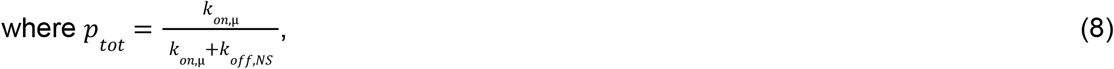

is the probability to bind the motif specifically when in the testing state, before dissociating macroscopically into the DNA dissociated free state. The key assumption, that the time spent in the testing state is short, collapses the two parameters *k*_off,NS_ and *k*_on,μ_ into a single parameter *p*_tot_, that only influences the macroscopic observables *k*_a_, *k*_d_ and *K*_D_ by how many times the TF needs to try and pass through the testing state before successful macroscopic association (state (1) → (3)) or dissociation (state (3) → (1)). As stated in^10^, we believe that this is a reasonable assumption, true for most search processes where a searcher is looking for a rare target among many false decoys. It is also straightforward to verify this assumption experimentally, by validating that the binding is specific. If the time spent in the testing state is non-negligible, the TF should bind nonspecific DNA without motifs with similar affinities compared to specific DNA with motifs. On the other hand, if the time spent in the testing state is negligible, the TF should bind non-motif DNA with much lower affinity than motif DNA. That KLF1 binds non-motif DNA with orders of magnitudes lower affinity than motif DNA (**Fig. S2A**) thus shows that the time spent in testing state must be very short.

For normal kinetic experiments (**Fig. 1H, middle**) *k*_off,u_ was estimated by fitting equation (4) to the association and dissociation rate constants for the motif mutation data. For the kinetic experiments where dissociation was performed with a high concentration of dark competitor (**Fig. 1H, middle**), *k*_off,u_ was estimated by taking the average of *k*_d_ for the motif mutation data. The three-state model predicts that the macroscopic dissociation rate *k*_d_ should approach *k*_off,u_ when blocking *k*_on,u_ with dark competitor, while assuming that *k*_off,NS_ is fast compared to *k*_off,u_.

### Derivations and equations for the target search model

The target search model (**Fig. 3H, Fig. S3H**) builds on the three-state model (**Fig. 1F**), but in this model the nonspecific “testing” state is broken up into many more microscopic states that describe TF search, including TF hopping and sliding during 1D diffusion along DNA. We define *k*_*on*,*max*_ times the free TF concentration as the association rate from solution to the nonspecifically DNA bound search mode, *p*_*bind*_ as the probability to bind the motif when being in the search mode and directly on top of the motif, and *k*_*off*,μ_ as the rate at which the TF exits the specifically bound state, to go into the search mode on top of the motif. We now define the model quantity *p*_*find*,*assoc*_ as the probability to find the motif given entering the search mode in an association process (appearing on a random position along the DNA) before dissociating from the DNA. *p*_*find*,*dissoc*_ is the probability to re-find the motif given entering the search mode in a dissociation process (appearing on top of the motif), before dissociating from the DNA. *p*_*find*,*assoc*_ and *p* _*find*,*dissoc*_ both depend on the microscopic model and parameters governing the search mode (more on this in the next section).

Below we will derive an equation to get ΔΔ*G* of motif binding for different flank mutations independent of *p* _*bind*_, *k* _*off*μ_ and *k* _*on*,*max*_, which are parameters that we believe do not change with flanking sequence variations. Namely, we will show that

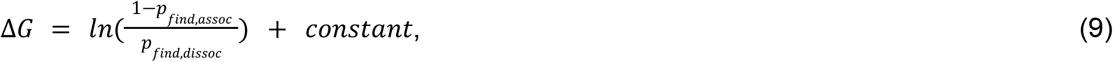

when we assume that *p* _*bind*_, *k* _*off*μ_ and *k* _*on*,*max*_ are constant across the considered set of sequences.

#### Proof

We start off by observing that, as for the three-state model, assuming that the time in the search mode is very short, equation (5) and (6) for the target search model becomes

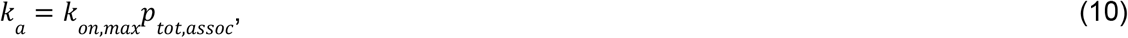

And

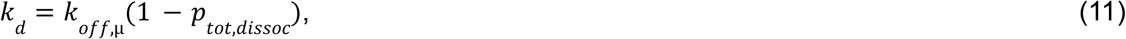

where *p*_tot,assoc_ is the probability to enter the specifically bound state on the motif (rather than just finding the motif as in *p* _*find*,*assoc*_) given entering the search mode in an attempted association process, before macroscopic dissociation, and *p*_tot,dissoc_ is the probability to re-enter the specifically bound state on the motif given entering the search mode in an attempted dissociation process, before macroscopic dissociation. Dividing equation (11) with (10) gives

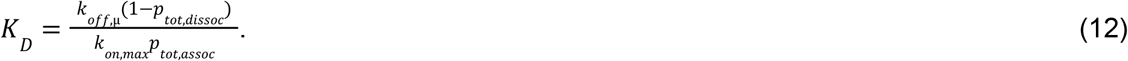

We will now derive equations for how *p* _*tot*,*assoc*_ and *p* _*tot*,*dissoc*_ depend on *p* _*find*,*assoc*_, *p* _*find*,*assoc*_ and *p*_*bind*_. We start by considering an attempted dissociation process (entering the search mode from the motif), in which the TF can re-enter the specifically bound confirmation on the motif by re-finding the motif and binding it (which includes scenarios in which the TF re-finds the motif, fails to bind, but re-finds the motif any number of times until the TF ultimately binds the motif successfully). The total probability of binding before macroscopic dissociation is thus

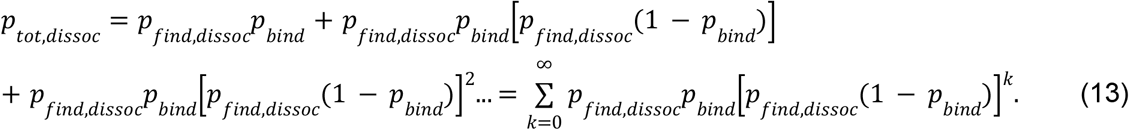

This is a geometric series, and since 0 < *p*_*find*,*dissoc*_< 1 and 0 < *p*_*bind*_ < 1, *p*_*find*,*dissoc*_(1 − *p*_*bind*_) < 1, this series converges to

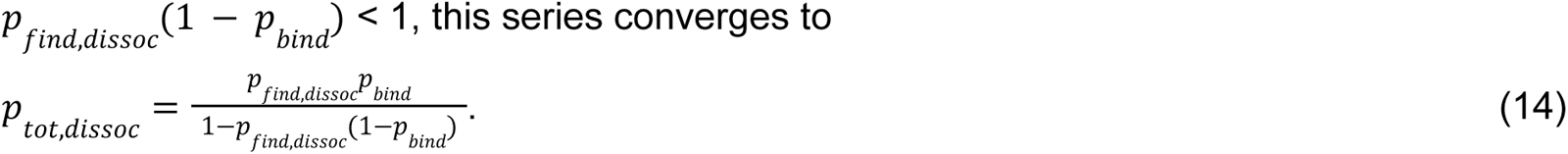

We now consider an attempted association process, entering the search mode on a random position along the DNA. The TF can find and enter the specifically bound confirmation on the motif by finding the motif and binding it (which includes scenarios in which the TF finds the motif and fails to bind it the first time, but re-finds the motif any number of times until the TF ultimately binds the motif successfully). The total probability of binding before macroscopic dissociation is thus

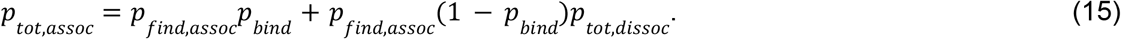

Inserting equation (14) into equation (15) gives us

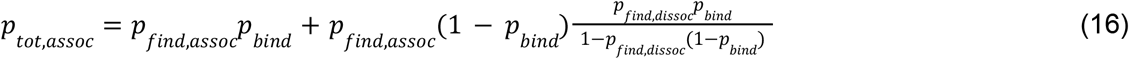

To relate to *K*_D_, we now insert equations (15) and (16) into equation (12), which after simplification gives

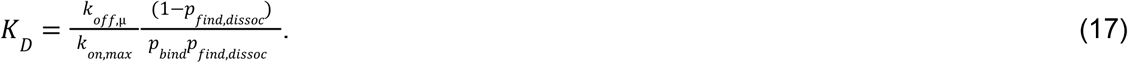

Equation (17) is a general result, true independent of the exact microscopic model and parameterization of the search mode (that govern *p* _*find*,*assoc*_ and *p* _*find*,*dissoc*_), and true also for sequences that have different *p* _*bind*_, *k* _*off*,μ_ or *k* _*on*,*max*_.

Finally we consider a set of DNA sequences that have the same *p* _*bind*_, *k* _*off*μ_ and *k* _*on*,*max*_, but different *p*_*find*,*dissoc*_ and *p* _*find*,*assoc*_. According to equation (17), their Δ*G* in *k*_B_*T* units can be calculated according to

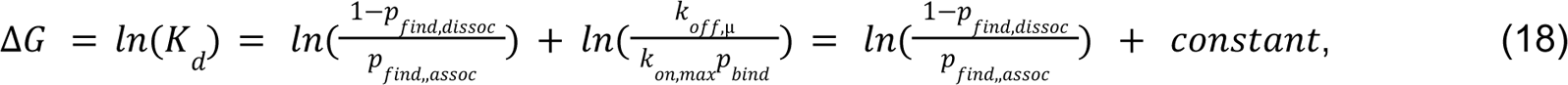

which is the sought equation (9) (QED).

### Simulations and parameterization of the target search model

To obtain *p*_*find*,*assoc*_and *p*_*find*,*dissoc*_, the target search process of the target search model (**Fig. 3H, Fig. S3H**) was evaluated using stochastic diffusion simulations on a discretized DNA with 1 bp steps, building on the simulations described in^9^. *p*_*find*,*assoc*_and *p*_*find*,*dissoc*_ both depend on the microscopic model parameters governing the search mode, which we define to include two forms of 1D diffusion: hopping and sliding. In our case, the microscopic parameters are: *k*_*hop*_, the rate of hopping to place the protein in a major groove an integer of DNA periods away from the previous position, *x*_*hop*_, the average hop length in bp, *p*_*flip*_, the probability that the TF flips

180 degrees and changes from sensing the forward or reverse strand given a hop, *D*^*helix*^, the rate at which the TF takes 1 bp helical diffusion step to the left or right, and *k*_*off*,*NS*_, the dissociation rate from the search mode and DNA into solution. *p* _*find*,*assoc*_ and *p* _*find*,*dissoc*_ can thus be viewed as a function of these microscopic parameters,

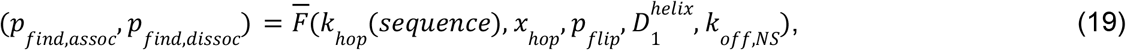

where *k*_*hop*_ (*sequence*) is a DNA sequence and/or position-dependent hop rate landscape (more on this below).

At any given simulation step when the TF is nonspecifically bound to the DNA (in the search mode), the TF can take a hop of average length *x* _*hop*_ to a nearby major groove with probability *k* /*p*, or take a 1 bp helical diffusion step with probability 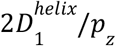 (to the left or right with equal probability), or dissociate macroscopically from the DNA with probability 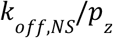,where

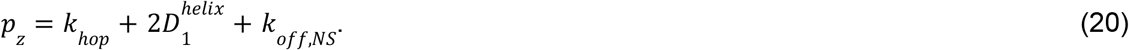

Hop lengths were sampled from a normal distribution with average 0 and standard deviation *x* _*hop*_, and then rounded off to the closest 10.5 bp to place the protein in the major groove an integer of DNA periods away from the previous position. Given a hop, the TF can flip to change if it is sensing the forward or reverse strand with probability *p* _*flip*_. Association and dissociation processes were sampled to estimate *p* _*find*,*assoc*_ and *p* _*find*,*dissoc*_, the probability to find the motif, i.e. be on top of the motif and sensing the forward DNA strand, before macroscopic dissociation. Association processes were initiated by sampling the starting position from a uniform distribution along the DNA, and equiprobable association to the forward or reverse strand. Dissociation processes were initiated with the TF on top of the motif on the forward strand, and the trajectory was then run until the TF found the motif again, or dissociated macroscopically.

The default values of model parameters were taken as 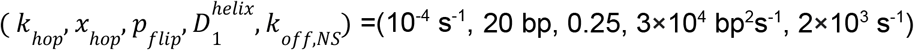 when not stated otherwise. These values were chosen to be within a factor of ∼2 of the values that have been determined experimentally for the *lac* repressor^7,9^, but while modifying them slightly to reflect that HaloTag-KLF1 DBD is smaller and perhaps a slightly weaker nonspecific binder than the *lac* repressor dimer.

In **Fig. 3I**, *k*_hop_ was modified on the forward strand at one specific position per simulation, while keeping all other hopping rates constant. When using a sequence dependent hop rate landscape (**Fig. 3J,K, Fig. S3I**), *k*_hop_ was calculated from the motif-likeness score at a given 9 bp sequence position according to

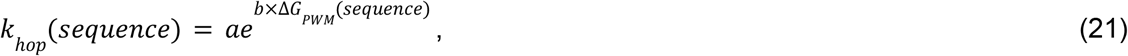

where the normalization parameter *a* is defined as

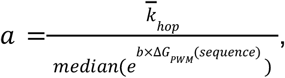

and 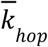 is the desired median *k*_hop_ given as an input parameter. Δ*G*_PWM_(*sequence*) was calculated from the motif PWM model fitted on our motif mutation data (**Fig. 1E**), and *b* was set to 0.1 to give a reasonable distribution of hop probabilities (*p* _*hop*_ = *k* _*hop*_ /*p* _*z*_), where higher values for *b* will give many hop probabilities very close to 0 or 1, and lower values for *b* will result in all hop probabilities being very similar.

To obtain ΔΔ*G* for different hop rate landscapes and model parameters, the *p* _*find*,*assoc*_ and *p* _*find*,*assoc*_ calculated by the above described simulations were inserted into equation (9).

### PWM models

The binding energy Δ*G*_PWM_ for a sequence in our PWM models is defined as

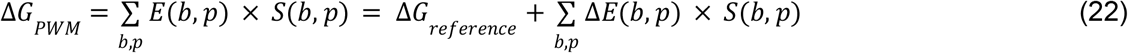

where *b* is the current base and *p* is the current position in the sequence. *S*(*b*,*p*) is a function that outputs 1 if the current sequence has the base *b* at position *p*, and otherwise outputs 0. *E*(*b*,*p*) is the absolute energy contribution of each base at each sequence position, and Δ*E*(*b*,*p*) is the corresponding relative energy contribution compared to some reference sequence with an experimentally determined binding energy of Δ*G*_reference_. A PWM model with a total of *P* sequence positions has 4×*P* parameters represented in the 2D matrix Δ*E*. The parameters of Δ*E* were fit by minimizing the mean squared deviation between the experimental and model predicted Δ*G*. The reference sequence was chosen so that Δ*G*_reference_ was the lowest energy in the training dataset, and the parameters in Δ*E* corresponding to this sequence were locked to 0. This reduces the number of free parameters to 3×*P*.

The input data for the flank PWM model consisted of 4,096 DNA sequences (174 bp in length), and their corresponding *in vitro* measured values for Δ*G* = *ln*(*K*_D_). These sequences all contain the consensus KLF1 binding motif CCACGCCA, and has a varying flanking sequence context obtained by both moving the motif position and mutating the flanking sequence with a fixed motif position. 10% of the data was then selected at random as test data, 10% as validation data, and the remaining 80% as training data. The model in **Fig. 2E-G** used the 10 bp flank sequences to the left and and right of the motif (20 sequence positions), resulting in 80 parameters whereof 60 were free in the fitting of Δ*E*(*b*,*p*).

The input data for training of the core motif PWM model (**Fig. 2E, top**) consisted of 356 sequences and their corresponding *in vitro* measured values for Δ*G*. These sequences contained all possible single, double, and a few higher order mutations to the consensus CCACGCCA motif. All sequences had the motif in the same position, in the same strong flanking context background. This model has a total of 36 parameters, whereof 27 were free in the fitting of Δ*E*(*b*,*p*).

### Genomic analysis of clustered motifs

TF-bound motifs in the human genome were acquired from a compiled dataset that measured DNase I footprints in 243 human cell and tissue types^40^. The footprints in this dataset were mapped to TFs using JASPAR PWM motifs^98^ and PWMScan^99^. PWMScan was first run over hg38, for all non-redundant human JASPAR motifs, with a p-value cutoff of 0.001. Looping over all PWMScan outputted TF motifs and DNase I footprints, a TF was mapped to a footprint if the center of the PWMScan motif overlapped with the footprint. The number of nearby same TF motifs was then calculated with the current JASPAR PWM motif with a p-value cutoff of 0.001, in a 100 bp window centered at the motif overlapping with the footprint. For each JASPAR TF, the average number of nearby motifs (y-axis on **Fig. S5A**) was then calculated by taking the average over all mapped footprints for this TF.

For null hypothesis testing, the number of nearby same TF motifs was also calculated when the bases in the 100 bp window were scrambled, while excluding the bases of the motif at the center from scrambling. Scrambling was performed by assigning a new position for each base by a randomized permutation. For a given TF-mapped footprint, this scrambling and counting of nearby motifs was performed 50 times, allowing for estimation of z-scores and p-values. For each JASPAR TF, this was performed for 200 TF mapped footprints per chromosome. For each JASPAR TF, the average number of nearby motifs when scrambling (x-axis on **Fig. S5A**) was then calculated by taking the average over all mapped footprints where scrambling was performed for this TF.

The average number of nearby motifs when scrambling was also calculated for each of the 50 bootstrapped iterations. This enabled calculation of z-scores (y-axis, **Fig. S5B**) with the equation

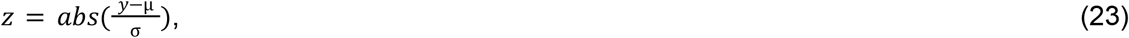

where *y* is the average number of nearby motifs when not scrambling, and µ and σ are the average and standard deviation of the bootstrap samples when scrambling.

### Deep learning model for flanks

The training, validation and test data for the deep learning flank model consisted of the same DNA sequences and measured *K*_D_ values as for the flank PWM model. The 174 bp sequences each contain the consensus KLF1 binding motif CCACGCCA. This motif was then cut out from the sequence along with the 30 closest nucleotides flanking each side of the motif. The resulting sequences were thus 69 base pairs long, centered on the binding motif. The values for *K*_D_ were normalized and logarithmized for the training and validation set, respectively.

PyTorch was used to implement the model with both the transformer and positional encoder class inheriting from torch.nn.Module. The model follows the classic transformer schema, starting off with an embedding layer, followed by a positional encoding layer. An encoder layer with 16 heads and a dropout of 0.2 makes up the transformer encoder. After this, a dropout layer is followed by a linear layer that outputs the values for ln(*K*_D_). In total, this model has 1,587,075 trainable and 8,960 untrainable parameters. The model was trained for 100 epochs, updating the trainable parameters to minimize the mean squared error between the model predicted and experimental ln(*K*_D_).

The positional encoding class uses the positional encoding (PE) functions from^100^ to create the tensors for odd and even positions. *pos* refers to the current position of the sequence being encoded. *i* refers to each dimension in the assigned positional encoder vector of length *d*.

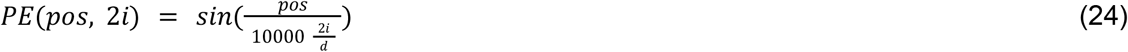

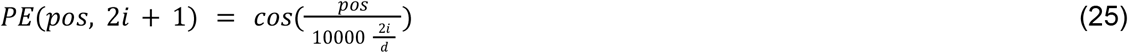

### Cell lines and cell culture

Cell culture was performed as described in^101^. Briefly, all experiments were performed in K562 cells (female, ATCC CCL-243). Cells were cultured in a controlled humidified incubator at 37°C and 5% CO2, in RPMI 1640 (Gibco, 11-875-119) media supplemented with 10% FBS (Omega Scientific, 20014T), and 1% Penicillin-Streptomycin-Glutamine (Gibco, 10378016).

Reporter cell lines were generated as in^102^. Briefly, reporter cell lines were generated by TALEN-mediated homology-directed repair to integrate donor constructs into the *AAVS1* locus by electroporation of 1 × 10^6^ K562 cells with 1 ug of reporter donor plasmid and 0.5 ug of each TALEN-L (Addgene no. 35431) and TALEN-R (Addgene no. 35432) plasmid. After 2 days, the cells were treated with 1,000 ng ml^−1^ puromycin antibiotic for 7 days to select for a population where the donor was stably integrated in the intended locus. These cell lines were not authenticated. All cell lines tested negative for mycoplasma.

Overexpression was performed by electroporation of 1 × 10^6^ K562 cells (selected to have desired constructs) with 2 ug of KLF1 overexpression plasmid (Addgene no. 120488) or 2 ug of pMAXGFP (Addgene no. 177825) as a control. Measurements were made 48 hours after electroporation.

### RT-qPCR and analysis

We harvested 1 million cells per replicate into buffer RLT (Qiagen) and homogenized the lysate with QIAshredder mini spin columns (Qiagen). We then purified RNA using the RNeasy Mini Kit (Qiagen), generate cDNA using the qScript Ultra SuperMix (Quantabio), and performed qPCR using SsoAdvanced Universal SYBR Green Supermix (BIO-RAD). We used primers against KLF1 from the overexpression plasmid (Addgene no. 120488) (TGCTCGTAGCGATGAACTTAC, CTGAAGGCACGTGGACATAA) and GAPDH (AGCACATCGCTCAGACAC, GCCCAATACGACCAAATCC) and calculated effects using the ΔΔCt method.

### SMF library design

For library design, background sequences were chosen from hg38 regions with high GC content (for good footprinting) that were not in K562 DNase peaks and did not contain simple repeats or key restriction enzyme sites. Motifs were then inserted with equal spacing and GCs immediately flanking the motif (consensus: 5’-a**gC**CAC**GC**CCA**gc**a-3’, broken: 5’-a**gC**CAT**GC**CCA**gc**a-3’) except for when explicitly varying the flanking sequence.

### Library cloning

Library cloning was performed as in^47^. Briefly, oligonucleotides of 350 bp in length were synthesized as pooled libraries (IDT), resuspended to 0.1 uM in TE buffer and diluted to 1 nM in H2O. Libraries were then PCR amplified to add on SMF primers (oJS114/oJS115) and gibson overhangs (cTF256, cTF257). Libraries were then cloned into a vector (pCL056^47^) with Gibson assembly using NEBuilder HiFi DNA Assembly Master Mix (NEB), transformed into Stable Competent E. coli (NEB, C3040), and isolated using QIAprep Spin Miniprep Kit (Qiagen, 27104).

### Methylation treatment

Methylation was performed as in^47^. Briefly, between 1-3 million cells were washed in ice cold PBS and then lysed on ice using the Omni-ATAC protocol. Isolated nuclei were then resuspended in 150 uL methylation mix that has been preheated to 37°C for 2 minutes: 100 uL cell methylation buffer (10% volume 10x M.CviPI reaction buffer (NEB, B0227S), 300 mM sucrose, 2.13 mM S-adenosylmethionine), and 50 uL of 4,000U/mL M.CviPI GpC methyltransferase (NEB, M0227L). Each methylation reaction was then incubated at 37 °C shaking at 1000 rpm for 7.5 min (unless specified otherwise) and quenched with 150 uL gDNA cell lysis buffer, 3 uL RNase A and 1 uL ProtK all from the Monarch gDNA extraction kit (NEB, T3010). We then extracted gDNA using the Monarch gDNA extraction kit, and quantified DNA Qubit using 1x dsDNA HS Assay Kit on a Qubit Flex Fluorometer.

### Enzymatic conversion

Enzymatic conversion was performed as in^47^. Briefly, methylated gDNA was digested using XbaI and AccI (NEB) to liberate a fragment containing the entire intact reporter sequence, and a 2-sided SPRI (0.4X followed by 1.8X) was performed to enrich for these fragments. DNA was then converted using the Enzymatic Methyl-seq conversion module (NEB).

### Amplicon library preparation and sequencing

Amplicon library preparation was performed as in^47^. Briefly, amplicon library preparation was performed in three rounds of PCR: 1) amplification out of the converted DNA with NEBNext Q5U Master Mix (NEB, M0597) (primers: cTF223, cTF218 - see Supp. Table 3) 2) addition of part of R1 and R2 sequencing primers with NEBNext Ultra Q5 Master Mix (primers: cTF401/402/, cTF403/404 - see Supp. Table 3) and 3) addition of remaining sequencing handles and sample indexes with NEBNext Ultra Q5 Master Mix (oID1-7, oID8-24 - see Supp. Table 3). Libraries were then quantified using Qubit 1x dsDNA HS Assay Kit for concentration and D1000 ScreenTape (Agilent Technologies, 5067-5582) for length. Libraries were pooled with 15% PhiX Sequencing Control v3 (Illumina, FC-110-3001) and 15% genomic EM-seq libraries for complexity and sequenced with 374 cycles on R1, 235 cycles on R2, 8 cycles on I1, and 4 cycles on I2 and with a MiSeq Reagent Kit v3 (600-cycle) (Ilumina, MS-102-3003) on a MiSeq.

### Genome-wide library preparation and sequencing

Genome-wide library preparation and sequencing was performed as in^47^. Briefly, methylated gDNA was sonicated to an average length of 400 bp using a Covaris E220 Focused-Ultrasonicator and combined with the methylated pUC19 and the unmethylated lambda-phage DNA (NEB). End prep and adaptor ligation was performed using the Enzymatic Methyl-seq kit (NEB). Conversion was performed identically to the amplicons. Libraries were constructed by PCR with the index primers for the EM-seq kit (NEB) with NEBNext Q5U Master Mix (NEB). Libraries were sequenced either by spiking into the amplicon libraries for complexity, or on their own on a Next-seq using standard primers with 2×36 bp reads.

### Genome-wide SMF analysis

Genome-wide SMF analysis was performed as in^47^. Briefly, reads were first trimmed to 36 bp using fastx_trimmer (http://hannonlab.cshl.edu/fastx_toolkit/index.html) and then aligned to the hg38, lambda, and pUC19 genomes using bwameth^103^. Duplicates were identified and removed using Picard MarkDuplicates (https://broadinstitute.github.io/picard/). Methylation fraction across the genome was called using MethylDackel (https://github.com/dpryan79/MethylDackel) and was aggregated either by sequence context or genomic position using custom scripts (available at https://github.com/GreenleafLab/amplicon-smf).

### Synthetic construct SMF analysis

Synthetic construct SMF analysis was performed as in^47^. Briefly, reads were aligned to a custom index (built from the individual PCR amplicons) using bwameth^103^. The resulting BAM files were filtered for alignment quality and uniqueness, unconverted reads were removed (by looking at the conversion of all non-GpC Cs), and bulk methylation was computed using MethylDackel (https://github.com/dpryan79/MethylDackel). Custom scripts were used to generate the single-molecule matrices and perform other quality control analyses. Library members were required to have at least 300 unique reads to be included in downstream analyses. A Snakemake pipeline with associated conda environments is available on GitHub (https://github.com/GreenleafLab/amplicon-smf) and via Zenodo (DOI: 10.5281/zenodo.13840888).

### Single-molecule state calling model

State calls were performed as in^47^ with a few modifications. Briefly, single-molecule states were assigned to each read using a maximum-likelihood approach. First, all possible single-molecule states (for a given amplicon) were enumerated (and precomputed to speed up future computations). A single-molecule state represents the occupancy status of each motif, as well as the position (start and end coordinates, snapped to the nearest GpC) of all nucleosomes covering the molecule. The full list of states was enumerated by taking the Cartesian product of the power set of the motifs (each can be bound or unbound) and all possible positions of as many nucleosomes will fit along the molecule (mandating that the nucleosomes start and end at GpCs, have at least one accessible GpC in a linker between any pair of two nucleosomes, have at least 25 bp in between bound TFs and nucleosomes, overlap the molecule with at least 30 bp, and are between 110-140 bp in length).

For each hypothetical state (TF occupancy and nucleosome positioning), the expected methylation signal was computed by determining whether each GpC is accessible, protected by a nucleosome, or protected by a TF. These GpC occupancy states were then converted into probabilities using different methylation probabilities (unobserved) for each of the three possible occupancy states. Methylation probabilities were decreased for short methylation times. This results in a matrix of size (number of states) x (number of GpCs) representing the probability that GpC j is methylated in state i. This matrix is then converted into a matrix *P* representing the probability of observing a converted base after the EM-seq reactions using the methylated (pUC19) and unmethylated (lambda) conversion rates from the controls. The maximum (log-)likelihood state for each input molecule (from single-molecule methylation matrix M, which is (number of GpCs) x (number of molecules)) was computed using a Bernoulli likelihood across all GpC positions in vectorized format, resulting in a vector of the most likely state for each input molecule. The code is available at https://github.com/GreenleafLab/klf_manuscript_code/classify_single_molecule_binding_v4.py.

### Partition function model

The partition function model was constructed as in^47^ with a few modifications. Briefly, the steady-state non-equilibrium thermodynamic model was constructed by deriving a function that assigns each molecule an energy. This energy function takes as input every possible single-molecule configuration from the state-calling model and counts the number of TFs (integers) and nucleosomes (including fractions of nucleosomes) occupying the enhancer. Each bound TF or nucleosome contributes a linear binding energy to the total energy of the molecule, except that nucleosomes bound to molecules with TFs also bound have their affinity reduced. Thus, three parameters (Δ*E*_TF_, Δ*E*_nuc_, Δ*E*_remodel_) are used to assign each molecule an energy. A given set of these parameters determines an energy, and therefore a Boltzmann probability, for each molecular state. The nucleosome energy was assigned to the value fit in^47^ due to the same locus being used. In **Fig. 5**, we then used maximum likelihood estimation to fit the two parameters (Δ*E*_TF_, Δ*E*_remodel_) that optimize the probability of seeing all observed molecules. The Simple Competition (**Fig. S5E-G**) was fit as above, although forcing the value of *E*_remodel_ to 0. In **Fig. 6H**, the now fixed parameters (Δ*E*_nuc_, Δ*E*_remodel_) are used, and the TF concentration (*[TF]*) is either fit directly or inferred (from value in **Fig. 5H**) to derive Δ*E*_TF_ from the PWM-predicted energies. The models were fit separately to each biological replicate of each experimental condition, and statistics were done on the independent fit values. Code is available at https://github.com/GreenleafLab/klf_manuscript_code/partition_function_model_functions.py and https://github.com/GreenleafLab/amplicon-smf/blob/master/workflow/scripts/fit_partition_function_model_v3.py.

In our implementation, Δ*E*_TF_ is defined by the relative probabilities of observing TF unbound versus bound DNA and is in units of k_B_T:

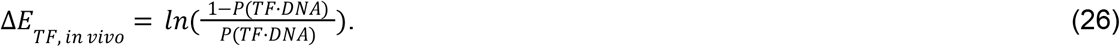

In comparison, *in vitro* measured energies are defined as:

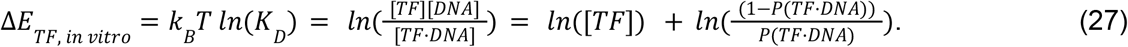

We can then (as in **Fig. 5H**) directly relate between these *in vitro* and *in vivo* defined energies as follows:

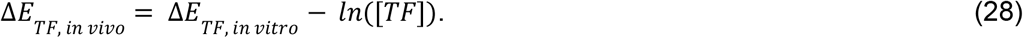

### Methylation and dissociation model

The kinetic model of methylation and dissociation was defined as a system of ordinary differential equations as follows,

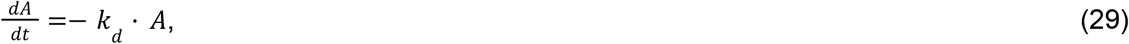

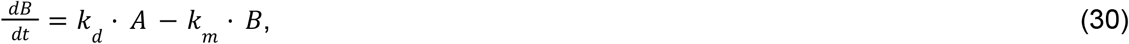

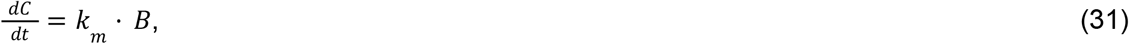

where the states *A, B*, and *C* refer to a TF-bound and unmethylated motif, an unbound and unmethylated motif, and an unbound and methylated motif, respectively, *k*_*d*_ is the TF dissociation rate, and *k*_*m*_ is the methylation rate.

To fit the model, *k*_*m*_ was input from a measured methylation rate (**Fig. 4H-I**) and A was assumed to be zero if the motif number was less than or equal to two and to linearly increase from three to seven motif numbers. Therefore, two parameters were fit across the average occupancy methylation time dataset: the initial t=0 average occupancy for the sequence with seven motifs (*A*, which was then scaled per motif number as described) and the dissociation rate, *k*_*d*_. Fitting was performed using scipy.optimize.curve_fit() and standard deviation on the parameter was calculated from the output covariance matrix, pcov, as follows: numpy.sqrt(numpy.diag(pcov)).

### RNA-seq data processing

RNA-seq dataset for K562s was sourced from ENCODE (experiment ENCSR000CPH) and the transcripts per million (TPM) for each KLF or Sp protein was identified. Code is available at https://github.com/GreenleafLab/klf_manuscript_code/RNA-seq_analysis.ipynb.

### BPNet predictions

BPNet predictions were performed as in^87^. Briefly, we used a BPNet model trained on KLF1 chromatin immunoprecipitation (ChIP) data in K562s (ENCODE: ENCSR550HCT). Each sequence from the *in vitro* KLF library in **Fig. 1A** are inserted at the center of one hundred 2114 bp random backgrounds. The average ChIP profile predictions were then obtained using the trained BPNet model (see https://github.com/kundajelab/bpnet/) and fold changes in ChIP profile predictions were calculated relative to the backgrounds input without the inserted sequence of interest.

## Data availability

All high-throughput sequencing for single-molecule footprinting datasets generated in this study are available through the NCBI Sequencing Read Archive (PRJNA1197470).

## Code availability

The synthetic SMF analysis and state-calling software are available on GitHub (https://github.com/GreenleafLab/amplicon-smf) and via Zenodo (DOI: 10.5281/zenodo.13840888). The RNA-seq processing scripts and the modified state-calling and partition function modelling functions for this manuscript are available on GitHub (https://github.com/GreenleafLab/klf_manuscript_code).

## Materials availability

This study did not generate new unique reagents.

## Additional Information

Supplementary Information is available for this paper. Correspondence and requests for materials should be addressed to Dr. William Greenleaf.

## Acknowledgements

We thank Eesha Sharma and Yuxi Ke for discussion on *in vitro* experiments, Vivekanandan Ramalingam for help with BPNet, Carl Nettelblad for discussions on *in vitro* deep learning model building, Abby Thurm, Polly Fordyce, and Lacra Bintu for feedback, Betty Liu for help with RNA-seq data, and Soon il Higashino for keeping our lab running. This work was supported by the Stanford Interdisciplinary Graduate Fellowship affiliated with Stanford Bio-X (J.M.S., B.R.D.), by grant numbers NSF GRFP DGE-1656518 (J.M.S., B.R.D., and M.M.H.) and Genetics/Dev Bio T32 training grant 1T32GM141828-01A1 (O.J.C.). E.M. acknowledges support from the Swedish Research Council (grants 2020-06459 and 2024-05105), the Foundation Blanceflor, Åke Wibergs Foundation, the Science for Life Laboratory (SciLifeLab), and computational resources provided by the National Academic Infrastructure for Supercomputing in Sweden (NAISS) partially funded by the Swedish Research Council through grant agreement no.

2022-06725. This work was supported by NIH grants UM1HG009436 and P50HG007735 (to W.J.G.). W.J.G. was a Chan Zuckerberg Biohub investigator and acknowledges grants 2017-174468 and 2018-182817 from the Chan Zuckerberg Initiative.

## Author Contributions

J.M.S., E.M., and W.J.G. conceived of this study. E.M. performed all *in vitro* experiments and data analysis. J.M.S performed and analyzed all *in vivo* experiments, with experimental assistance from B.R.D. and M.M.H. and analysis assistance from B.R.D. E.M. and J.M.S. derived models and equations, and performed simulations. T.F. implemented the flanking sequence machine learning model. O.J.C. performed BPNet *in silico* experiments. J.M.S., E.M., and W.J.G. wrote the manuscript, with input from all authors. E.M. and W.J.G. jointly supervised the work.

## Supplementary Information

<u>Document S1</u>

Figures S1–S6

<u>Table S1</u>

Library DNA sequences and annotations for HiTS-FLIP experiments. Fluorescence values, and fit *k*_a_/*k*_a,ref_, *k*_d_ and *K*_D_ for kinetic and equilibrium experiments.

<u>Table S2</u>

Oligo libraries installed at the reporter for SMF experiments.

<u>Table S3</u>

DNA sequences used for primers in targeted amplification for SMF library sequencing and SMF library cloning. Primers and oligos used for HiTS-FLIP library sequencing and binding measurements. Translated protein amino acid sequence for KLF1 DBD purification.

<u>Table S4</u>

Summary metrics (e.g. average TF occupancy) for all SMF experiments.

<u>Table S5</u>

Single-molecule state model calls for all SMF experiments.

<u>Table S6</u>

Sample information for all SMF experiments.

---

**Figure S1.**
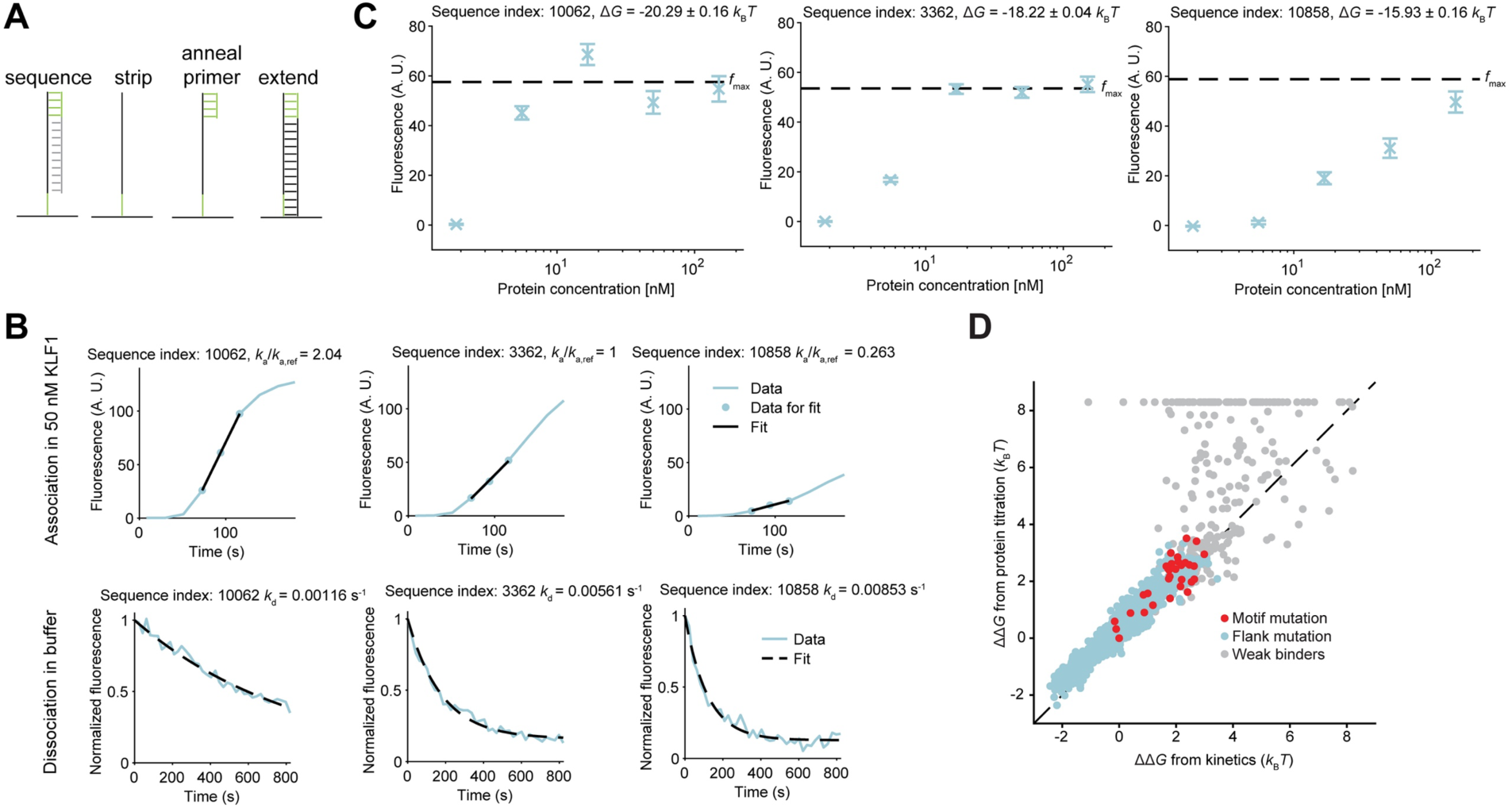
A) Schematic of preparation of the HiTS-FLIP array. B) Example association (top) and dissociation (bottom) curves, and curve fits used to quantify relative association
rates (*k*_a_/*k*_a,ref_) and dissociation rates (*k*_d_), for three flank mutations, all with a strong consensus motif (CCACGCCCA). C) Example binding curves when titrating the KLF1 DBD concentration, and quantified binding energies (Δ*G* = ln(*K*_D_), for the same flank mutations as in B. Dotted line represents the fluorescence signal at saturating KLF1 DBD concentration (f_max_) for each sequence in the titration experiment. D) Relative binding energies (ΔΔ*G*, relative to sequence index 3362 in panels B and C) obtained via the protein titration experiment and the kinetic experiment.

**Figure S2.**
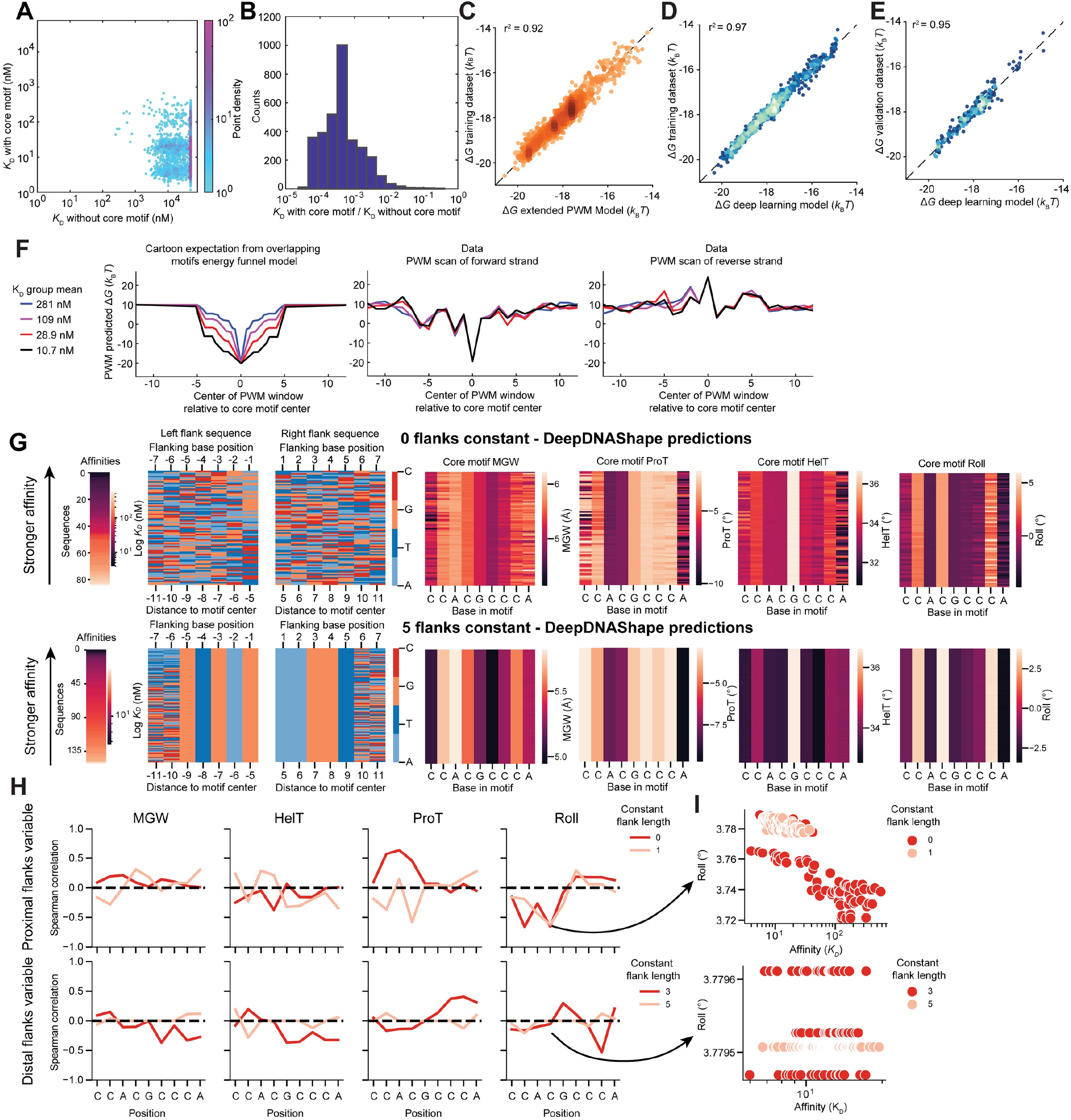
A) Relationship between binding affinities with and without the core motif present for different flanking sequence context variations. For the sequences without core motif, the motif sequence has been mutated from CCACGCCA to TACTTAGAT. B) Histogram of the ratio between binding affinities with and without the core motif. C) Relationship between extended Position Weight Matrix (PWM) model predicted and measured binding energies on the training dataset. D-E) Relationship between machine learning model predictions and measured binding energies for the training (D) and validation (E) datasets. F) Average core motif PWM scores (Δ*G*_PWM_) sliding the window across sequences for each affinity group (colors) in the forward stand (middle), the reverse strand (right), and as a cartoon representation of results if flanking sequence effects were driven by low affinity sites overlapping the core motif. The selected sequences all contain the consensus CCACGCCCA motif, and the entire flank is varied by changing the motif position (flank constant length 0). G) DeepDNAshape predictions across variable sequence contexts, sorted by affinity, with zero (top) or five (bottom) constant flanking base pairs. Shape predictions of minor groove width, propeller twist, helical twist, and roll are across the core motif when including the seven flanking base pairs on each side. The sequence content of variable flanking regions are represented on the left. H) Correlation coefficients between shape parameters at specific motif positions and measured affinities for when proximal flanks (top) or distal flanks (bottom) are varied. I) Example relationships between affinity and Roll at the fourth base in the core motif to demonstrate the negligible range in Roll across proximal flank variations and the lack of correlation across distal flank variations.

**Figure S3.**
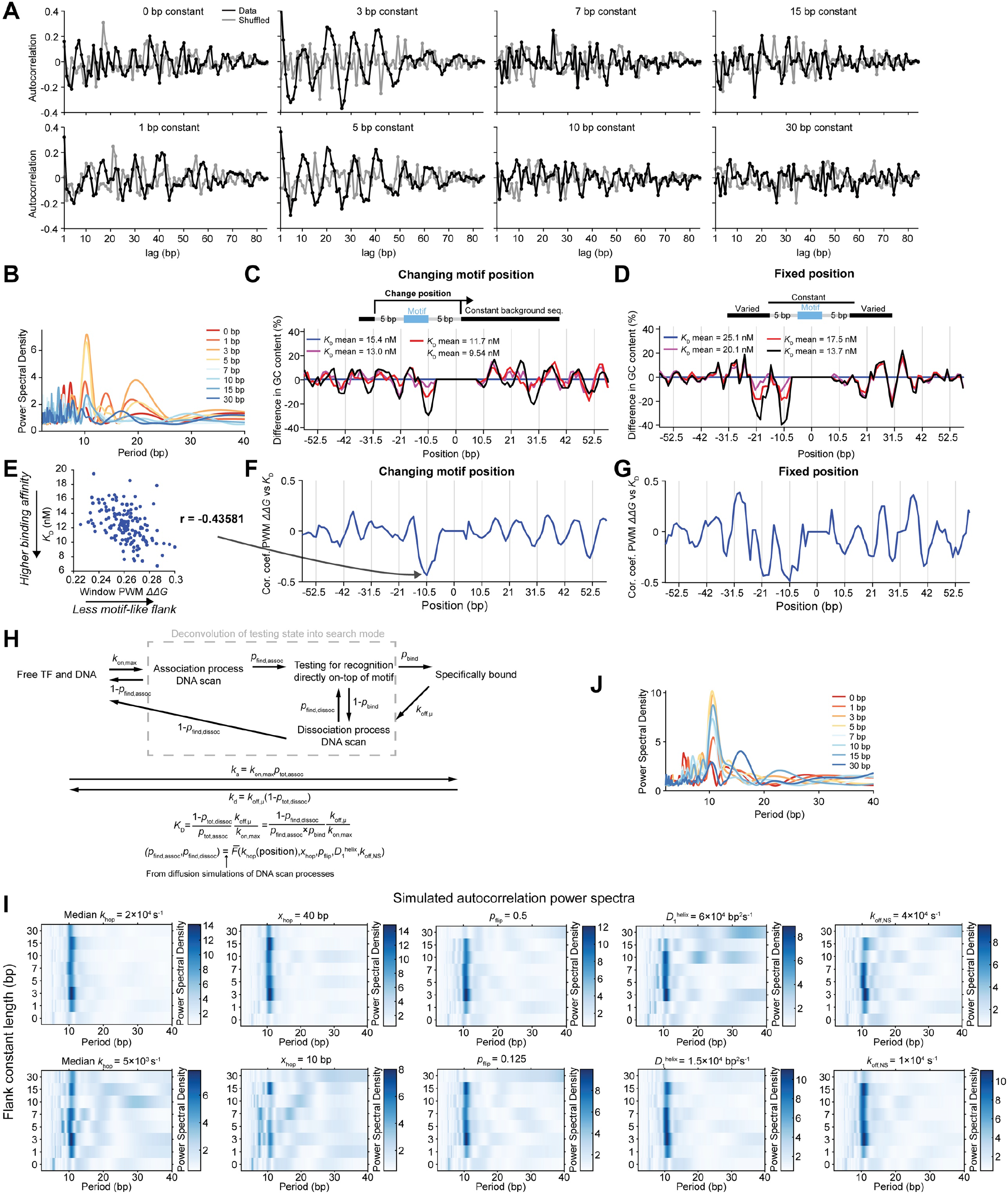
A) Autocorrelation of the binding energies across motif positions with variable lengths of constant flanks (black) and shuffled data (gray). B) Power spectra of the autocorrelations in A where the hue denotes the constant flank length. Contains the same data as in **Fig. 3D**, but with the different flank lengths plotted as different curves. C-D) The difference in GC content across flanking positions relative to the core motif for different affinity groups (colors) compared to the weakest affinity group (blue) for both the changing (left) and fixed (right) motif position libraries. E) Relationship between the window PWM score one helical turn to the left from the motif and measured affinity. The window PWM score is calculated as the average Δ*G*_PWM_ in a 4 bp window, centered at the current sequence position, over both the forward and reverse strand. F-G) The correlation coefficient between the window PWM score at variable positions away from the motif and measured affinity for both the changing (left) and fixed (right) motif position libraries. H) Schematic of target search model. I) Simulated autocorrelation power spectra as in **Fig. 3K** for 4-fold perturbations of the model parameters, with default values (in Fig. 3K) 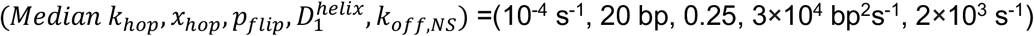. J) Power spectra as in B for simulated autocorrelations with the default parameter values. Contains the same data as in **Fig. 3K**, but with the different flank lengths plotted as different curves.

**Figure S4.**
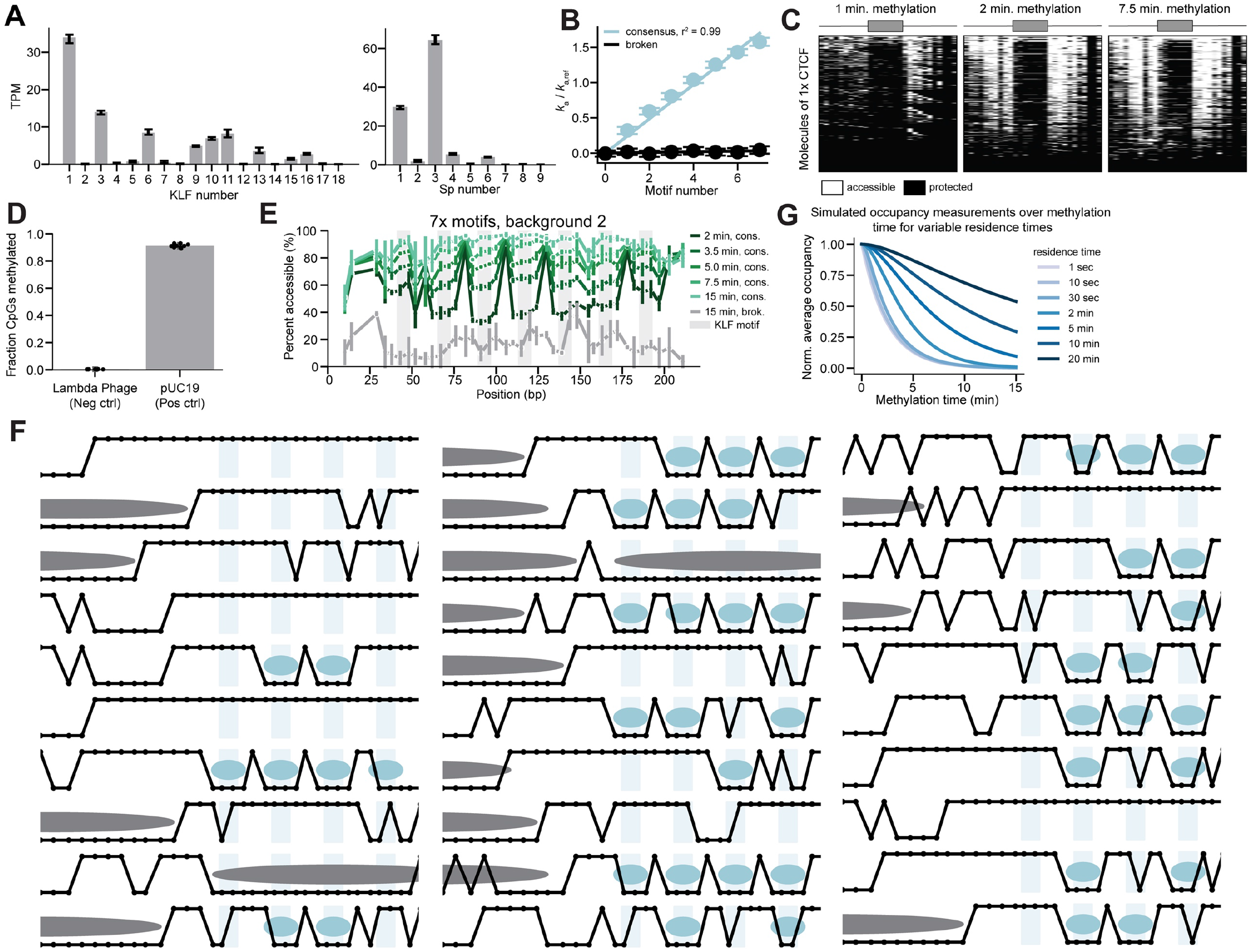
A) Transcripts per million from RNA-seq data on K562 cells across KLF (left) and Sp (right) factors. B) Relationship between motif number and *in vitro* relative association rate. Line is a linear fit. Error bars are bootstrapped s.d. C) Summary plots of single molecules as in Fig. 4E for constructs with one high affinity CTCF motif at variable methylation times. D) Methylation probabilities at CpGs from the fully methylated (pUC19) and fully unmethylated (Lambda phage) EM-seq controls. E) Average methylation for the seven consensus (greens) and broken (gray) motifs across methylation times. Error bars represent s.e.m. from two biological replicates. F) Exemplar data for 30 random single molecules, each of four consensus motifs background two at two minute methylation, and binding model calls for nucleosomes (gray) and KLF (blue). For each molecule, points on the top are methylated (accessible) and points on the bottom are unmethylated (inaccessible). G) Simulated TF occupancy as measured by SMF across a methylation time course for variable TF residence times.

**Figure S5.**
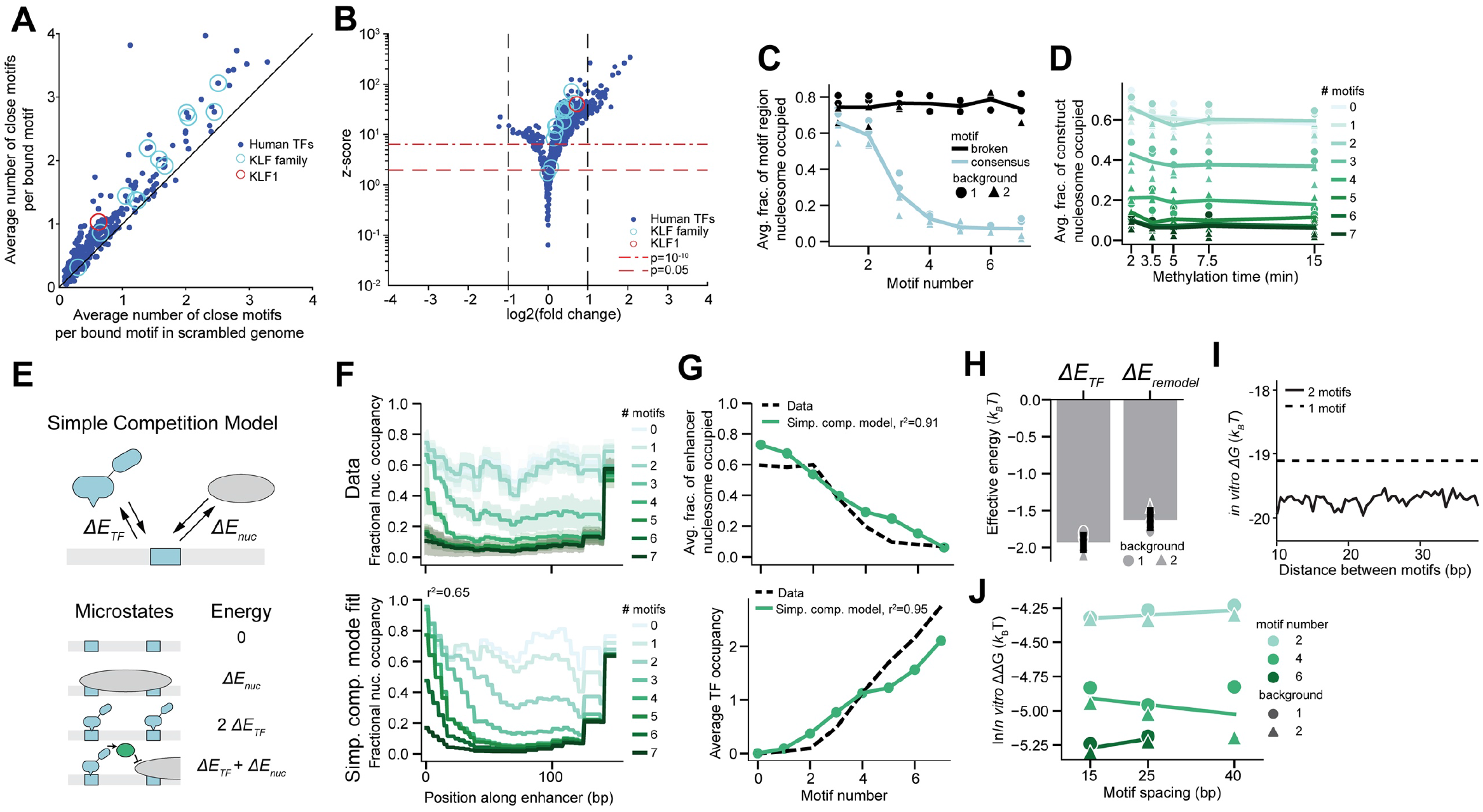
A) The average number of nearby motifs (closer than 50 bp, identified by JASPAR) per bound motif (identified by DNAse-seq) across transcription factors relative to if the sequence region around the bound motif is scrambled. Blue circles denote KLF family members and the red circle denotes KLF1. B) The relationship between log fold changes and z-scores for data in A, where fold change is the ratio between average number of nearby motifs and average number of nearby motifs when scrambling. C) The relationship between motif number and the average fraction of the motif-containing region of constructs that is nucleosome occupied (rather than fixed end-to-end distance across constructs) across two backgrounds (symbols) and two biological replicates at 7.5 minute methylation. D) The relationship between methylation time and the average fraction of constructs that are nucleosome occupied across motif number (color) and for two background contexts (symbols) and two biological replicates. E) Schematic of the Nucleosome Destabilization model describing competition of nucleosomes and TFs for binding to DNA F) The fractional nucleosome occupancy across the constructs with 0-7 consensus motifs for measured (top, 7.5 minutes of methylation) and the fit Simple Competition model (bottom). Error bars are standard deviations across two backgrounds and two biological replicates. G) The relationship between motif number and nucleosome occupancy (top) or average TF occupancy (bottom) for measured data (black) and the fit Simple Competition model (green). H) Best-fit parameters for KLF binding using the Nucleosome Destabilization model. Error bars are s.d. Across two backgrounds and two biological replicates. I) *In vitro* binding energy when changing the distance between two consensus motifs. Average one motif binding energy denoted with dashed line. J) The relationship between *in vitro* binding energy and motif spacing in the SMF library for two backgrounds.

**Figure S6.**
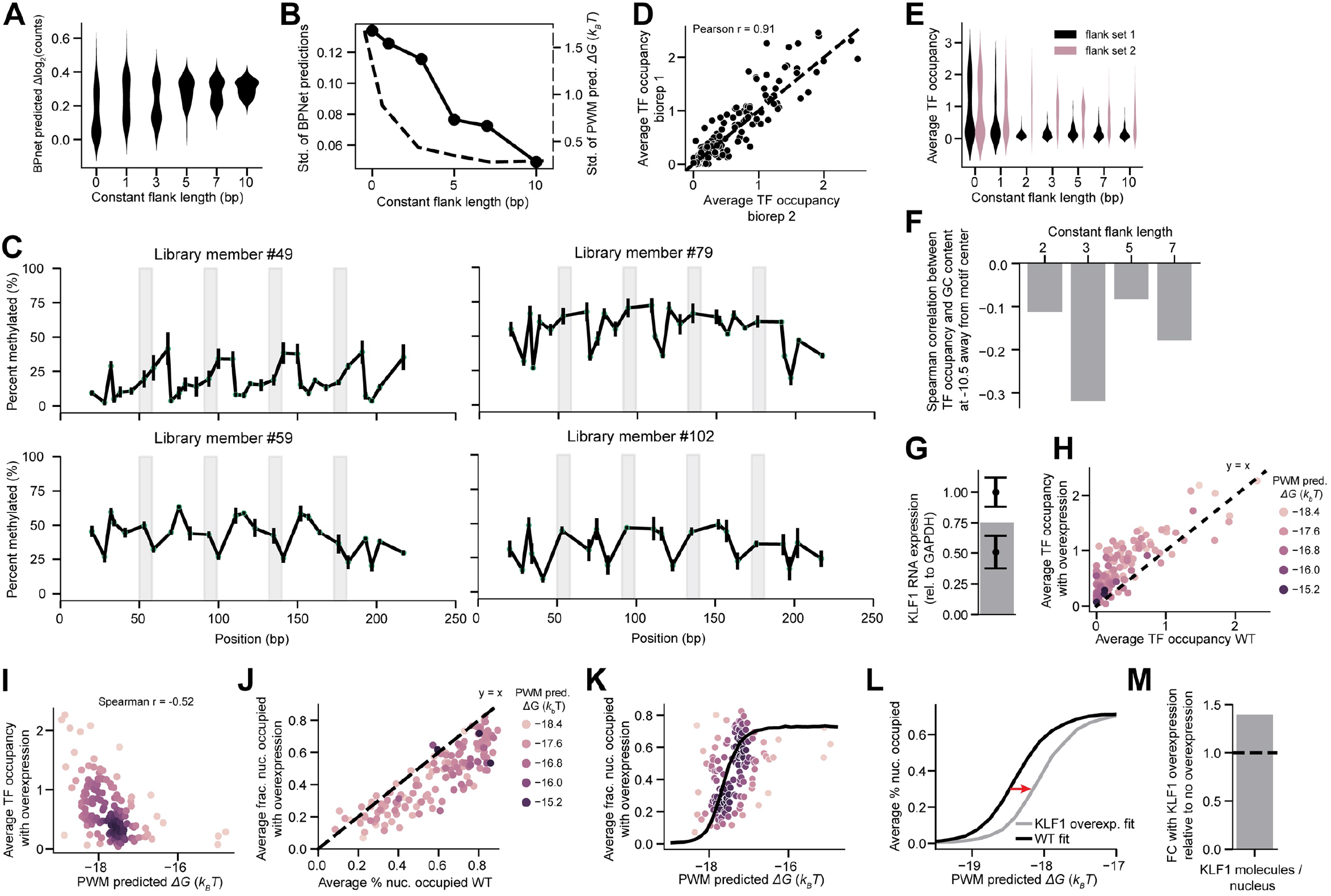
A) The distribution of log fold-changes of BPNet predicted KLF1 chromatin immunoprecipitation read counts for flanking sequence variations in **Fig. 6A** across increasing lengths of constant flanks. B) Relationship between the length of constant flanking sequence and the standard deviation of both BPNet-predictions in A (solid) and *in vitro* PWM-predicted energies (dashed). C) Representative examples of non-KLF binding (occupancy outside of motifs) to the library in **Fig. 6B** that is visible in the average methylation signal. Error bars represent s.e.m. from two biological replicates. D) Reproducibility in average TF occupancy measurements (2 minute methylation) for the library in **Fig. 6B** between two biological replicates. E) The distribution of TF occupancy measurements (2 minute methylation) across increasing constant flank lengths for two flank sets and two biological replicates. F) Spearman correlations between average TF occupancy and GC content one helical turn away from the motif center across constant flank lengths for flank set two (two biological replicates, p values = 0.3, 0.0007, 0.4, 0.05). G) Quantification of transient KLF1 overexpression by RT-qPCR for two biological replicates. Error bars are s.d. Over three technical replicates. H) Average TF occupancy (2 minute methylation) with KLF1 overexpression relative to no overexpression. Colors denote PWM predicted binding energies. I) The relationship between PWM predicted binding energies and average TF occupancy (2 minute methylation) with KLF1 overexpression. Color represents density. J) Comparison of the fraction of molecules that are nucleosome occupied (7.5 minute methylation) as in G. K) The relationship between PWM predicted energies and the fraction of molecules that are nucleosome occupied. Color represents density. Line is the fit two parameter model described in **Fig. 6C**. L) Comparison of two parameter model fits with (gray) and without (black) KLF1 overexpression. M) Fit KLF concentration from KLF1 overexpression in J relative to fit without overexpression.

